# Highly sensitive and specific multiplex antibody assays to quantify immunoglobulins M, A and G against SARS-CoV-2 antigens

**DOI:** 10.1101/2020.06.11.147363

**Authors:** Carlota Dobaño, Marta Vidal, Rebeca Santano, Alfons Jiménez, Jordi Chi, Diana Barrios, Gemma Ruiz-Olalla, Natalia Rodrigo Melero, Carlo Carolis, Daniel Parras, Pau Serra, Paula Martínez de Aguirre, Francisco Carmona-Torre, Gabriel Reina, Pere Santamaria, Alfredo Mayor, Alberto García-Basteiro, Luis Izquierdo, Ruth Aguilar, Gemma Moncunill

## Abstract

Reliable serological tests are required to determine the prevalence of antibodies against SARS-CoV-2 antigens and to characterise immunity to the disease in order to address key knowledge gaps in the context of the COVID-19 pandemic. Quantitative suspension array technology (qSAT) assays based on the xMAP Luminex platform overcome the limitations of rapid diagnostic tests and ELISA with their higher precision, dynamic range, throughput, miniaturization, cost-efficacy and multiplexing capacity. We developed three qSAT assays to detect IgM, IgA and IgG to a panel of eight SARS-CoV-2 antigens including spike (S), nucleoprotein (N) and membrane (M) protein constructs. The assays were optimized to minimize processing time and maximize signal to noise ratio. We evaluated the performance of the assays using 128 plasmas obtained before the COVID-19 pandemic (negative controls) and 115 plasmas from individuals with SARS-CoV-2 diagnosis (positive controls), of whom 8 were asymptomatic, 58 had mild symptoms and 49 were hospitalized. Pre-existing IgG antibodies recognizing N, M and S2 proteins were detected in negative controls suggestive of cross-reactive to common cold coronaviruses. The best performing antibody isotype/antigen signatures had specificities of 100% and sensitivities of 94.94% at ≥14 days since the onset of symptoms and 96.08% at ≥21 days since the onset of symptoms, with AUC of 0.992 and 0.999, respectively. Combining multiple antibody markers as assessed by qSAT assays has the highest efficiency, breadth and versatility to accurately detect low-level antibody responses for obtaining reliable data on prevalence of exposure to novel pathogens in a population. Our assays will allow gaining insights into antibody correlates of immunity required for vaccine development to combat pandemics like the COVID-19.

## INTRODUCTION

In a globalized world where emerging infectious diseases of broad distribution can put at stake the health and economy of millions of people, there is a need for versatile and reliable serological tools that can be readily applicable (i) to determine the seroprevalence of antibodies against any new pathogen and, more importantly, (ii) to characterise immunity to the disease at the individual and community levels. In the case of the COVID-19 pandemic caused by SARS-CoV-2, one of the main priorities since the beginning of the epidemics in China by the end of 2019 (1) was to ascertain the percentage of the population that had been exposed to the virus, considering that a considerable number of people could have been asymptomatic (2)(3). The lack of sensitive and specific serological tests early in the COVID-19 pandemic delayed the precise estimation of the burden of infection for the rational implementation of public health measures to control viral spread (4). Furthermore, immunological assays that can measure a high breadth of antibody types and specificities are needed to dissect which are the naturally acquired protective responses and identify correlates of immunity (5). Additionally, when a vaccine becomes available, such assays would be valuable to evaluate immunogenicity of candidate vaccines and monitor duration of immunity at the population level (6).

Common tools for antibody studies are (i) rapid diagnostic tests (RDT) as point of care (POC) devices that usually measure either total immunoglobulins or IgG and IgM, qualitatively (7), (ii) traditional enzyme-linked immunosorbent assays (ELISA) (8) that can quantify different isotypes and subclasses of antibodies against single antigens at a time, and that require certain previous expertise, personnel and equipment, and (iii) chemiluminescent assays (CLIA), widely used in clinical practice, faster and with higher throughput than ELISA (9). The performance characteristics of the commercial kits available in the early months of the COVID-19 pandemic were questionable (10), while external evaluations validating their reliability and accuracy were not published. A number of in-house ELISA assays have also been developed in hospital and research laboratories (11), but they have the limitations that (i) a relatively large amount of sample is required, (ii) the large surface area of the individual microplate wells and the hydrophobic binding of capture antibody can lead to non-specific binding and increased background, and (iii) most ELISAs rely upon enzyme-mediated amplification of signal in order to achieve reasonable sensitivity (12).

An alternative technique that offers the benefits of ELISA but also a larger dynamic range of antibody quantification and higher sensitivity (12)(13) is based on the xMAP Luminex® platform (www.luminexcorp.com/bibliography). Secondary antibodies are labelled with fluorescent phycoerythrin (PE) directly or with biotin that mediates binding to streptavidin-R-phycoerythrin (SAPE), which does not depend on an additional reaction. The technique has the added value of higher throughput (up to 384-well plate format), increased flexibility, and lower cost with the same workflow as ELISA, particularly if using magnetic MagPlex® microspheres. Paramagnetic beads allow for automation of workflow and better reproducibility compared to the previous generation of MicroPlex® microspheres. Since the beads have the capture antigen immobilized on their much smaller surface area compared to a 96-well microplate well, reduced sample volumes are required and non-specific binding is diminished (14). Furthermore, a chief advantage over ELISA is the multiplex nature of the assay that allows measuring antibodies to different antigens simultaneously. This increases the probabilities to detect a positive antibody response due to the heterogeneity of the human response and therefore it has a higher sensitivity relevant for identifying seropositive individuals. The Luminex technology, capable of measuring simultaneously antibodies against 50 (MAGPIX®), 80 (Luminex 100/200®) and up to 500 different antigens (FlexMap3D®), makes it an invaluable tool for antigen and epitope screening. Finally, its versatility to set up adapted antigen panels makes Luminex an excellent platform to ensure better preparedness for faster response to future emerging diseases and pandemics.

Here we report on the establishment and validation of three quantitative suspension array technology (qSAT) assays to measure IgM, IgA and IgG antibodies against eight SARS-CoV-2 antigens, based on the adaptation of previous in-house protocols that measured antibodies to other infectious diseases, including malaria (15)(16)(17)(18). Due to the need to process a large amount of samples with the minimal time and cost during a pandemic like the COVID-19, we optimized several conditions to reduce the duration of the assays and report them here for the three main isotypes that have proved useful for seroprevalence studies (19).

## METHODS

### Samples

Positive samples were 115 plasmas from individuals with a confirmed past/current diagnosis of COVID-19. One hundred and eleven had SARS-CoV-2 infection confirmed by real time reverse-transcriptase polymerase chain reaction (rRT-PCR). Fifty-five were recruited in a study of health care workers in Hospital Clínic in Barcelona, most of them with mild symptoms, 1 of them hospitalized and 6 without symptoms, all rRT-PCR positive (19). Fifty-seven were COVID-19 patients recruited at the Clínica Universidad de Navarra in Pamplona (Spain), of which 48 had severe symptoms and were hospitalized and 9 had mild symptoms (one clinically diagnosed with positive radiology and serology, and negative rRT-PCR); 3 were asymptomatic health workers with positive diagnosis confirmed by four serological tests but no rRT-PCR data. Time since onset of symptoms ranged from 0 to 46 days. Positive samples were used individually or as pools of up to 20 samples depending on the tests. For optimization tests, only a subset of samples were used. Negative controls were plasmas from 128 healthy European donors collected before the COVID-19 pandemic, and were used individually. Numbers of positive and negative samples were in line with protocol recommendations from the Foundation for Innovative New Diagnostics (FIND). *Ethics.* Samples analyzed in this study received ethical clearance for immunological evaluation and/or inclusion as controls in immunoassays, and the protocols and informed consent forms were approved by the Institutional Review Board (IRB) at HCB (Refs. CEIC-7455 and HCB/2020/0336) or Universidad de Navarra (Ref. UN/2020/067) prior to study implementation.

### Antigens

The Receptor-Binding Domain (RBD) of the spike (S) glycoprotein of SARS-CoV-2, the leading vaccine candidate target, was selected as the primary antigen to develop the initial qSAT assay because (i) S is one of the most immunogenic surface proteins together with the nucleocapsid protein (N) (20) (ii) RBD is the fragment of the virus that mediates binding to the host receptor ACE2 in the lung cells (21) (iii) antibodies to RBD correlate with neutralizing antibodies (20)(22) that could be associated with protection based on studies of other coronaviruses and animal models (23–26), and (iv) an ELISA based on this same protein has received FDA approval for COVID-19 serology (11). The RBD was from the Krammer lab (Mount Sinai, New York, USA) (11) and the S antigen was produced in-house using Chinese Hamster Ovary (CHO) cells transiently-transfected with the Krammer plasmid followed by purification of the recombinant protein from 4-day culture supernatants using nickel affinity chromatography. The multiplex antigen panel was completed with commercial S1 (GenScript Biotech, Netherlands) and S2 (Sino Biologicals, Germany) proteins, and in-house produced nucleocapsid (N) and membrane (M) recombinant proteins. *Escherichia coli* codon optimized versions of full-length N and M antigens were cloned at ISGlobal into a pET22b expression vector, fusing an in-frame C-terminal 6xHis-tag. Recombinant N and M proteins were expressed in *E. coli* BL21 DE3 by pET22b-N and pET22b-M transformation and induction with 0.5 mM isopropyl-β-d-thiogalactopyranoside (IPTG) when OD_600_ reached 0.6-0.8, followed by 5 h incubation at 37°C or 25°C, respectively. Bacterial pellets were resuspended in binding buffer containing 20 mM sodium phosphate, 0.5 M NaCl, 20 mM imidazole, 0.2 mg/mL Lysozyme, 20 µg/mL DNAse, 1 mM PMSF and 1 mM MgCl_2_, and lysed by sonication. Lysates were centrifuged at 14,000 rpm and proteins purified by affinity chromatography using a Ni^2+^ column (1 mL GE Healthcare HisTrap HP) and imidazole gradient elution in an AKTA Start protein purification system. M and N proteins were concentrated and buffer changed to phosphate buffered saline (PBS) using Microcon-10 KDa centrifugal filter units (Millipore). For N-terminal (residues from 43 to 180) and C-terminal fragments (residues from 250 to 360) of N, two constructs were designed at CRG depending on secondary structure predictions. The encoding sequences were synthesized and inserted into a plasmid pETM14 with the N-terminal 6xHis-tag under the control of a T7 promoter, and recombinant plasmids transformed into *E. coli* BL21 DE3 competent cells. Briefly, *E. coli* containing the plasmid was grown and the protein expression was induced by addition of IPTG 0.2 mM for 16 h at 18°C. Pelleted cells were resuspended in Buffer A (Tris 20 mM, 250mM NaCl, 10mM Imidazole) supplemented with 0.5% Triton-X100 Substitute (Sigma) and complete mini EDTA-free protease (Roche), sonicated, and centrifuged (30 min, 4°C, 30000 g). The N-terminal, and the C-terminal recombinant proteins containing a N-terminal 6xHis-tag were purified from the resulting supernatant using Hitrap Ni-NTA column (GE Healthcare, Uppsala, Sweden) according to the manufacturer instructions. After washing with Buffer A, the antigen was eluted using linear gradient with buffer B (Buffer A supplemented with 500 mM Imidazol). The fractions of interest were dialyzed against PBS 1x and concentrated by Vivaspin 5 KD (Millipore, France). The antigens produced were quantified using a bicinchoninic acid (BCA) protein assay kit (Pierce) and their purity controlled by Sodium Dodecyl Sulfate - PolyAcrylamide Gel Electrophoresis (SDS-PAGE).

### Antigen coupling to microspheres

Different test concentrations of protein antigens were coupled to magnetic MAGPLEX 6.5 µm COOH-microspheres from Luminex Corporation (Austin, TX) in reactions of a maximum of 625,000 beads, at 10,000 beads/µl (15). First, beads were washed twice with 62.5 µl of distilled water using a magnetic separator (Life Technologies, 12321d), and resuspended in 80 µl of activation buffer, 100 mM monobasic sodium phosphate (Sigma, S2554), pH 6.2. To activate the beads for cross-linking to proteins, 10 µl of 50 mg/mL sulfo-N-hydroxysulfo-succinimide (Thermo Fisher Scientific, 24525) and 50 µL of 50 mg/mL 1-ethyl-3-[3-dimethyl-aminopropyl]-carbodiimidehydrochloride (Thermo Fisher Scientific, 22981) were simultaneously added to the reaction tubes, mixed and incubated at room temperature (RT) for 20 min in a rotatory shaker and protected from light. Next, beads were washed twice with 62.5 µl 50 mM morpholineethane sulfonic acid (MES) (Sigma, M1317) pH 5.0, in a 10,000 beads/ µl concentration. After beads activation, antigen were added to the reaction tubes at three different concentrations (10, 30 and 50 μg/mL) and left at 4°C overnight (ON) on a rotatory shaker protected from light. On the following day, coupled-beads were brought to RT for 20 min in agitation, and blocked by incubating them with 62.5 µl PBS (Sigma) + 1% bovine serum albumin (BSA, Biowest) + 0.05% sodium azide (Sigma, S8032) (PBS-BN) in agitation during 30 min at RT and protected from light. Beads were washed twice with PBS-BN using the magnetic separator. To determine the percentage recovery of beads after the coupling procedure, coupled beads were resuspended in 62.5 µL PBS-BN and counted on a Guava PCA desktop cytometer (Guava Technologies, Automated cell counter, PC550IG-C4C/0746-2747). In all washing and resuspension steps, beads were softly vortexed and sonicated for 30 sec. Antigen-coupled beads were validated by incubating them with serial dilutions of anti-histidine-Biotin antibody for antigens with a histidine-tag (Abcam, ab27025). To choose an appropriate coupling concentration, IgG and IgM levels were measured in 11 serial dilutions 3-fold of a pool of 20 positive samples and titration curves compared for each protein. Coupled beads were stored multiplexed at 2000 beads/µL PBS-BN at 4°C and protected from light until use.

### Incubation of samples with antigen-coupled microspheres

We compared the performance of the assays when a subset of positive and negative plasma samples were incubated at different dilutions with the antigen-beads for 1 h or 2 h at RT in relation to our previous protocol ON at 4°C. Antigen-coupled beads, initially including RBD singleplex, were added to a 96-well µClear® flat bottom plate (Greiner Bio-One, 655096) at 2000 beads/well in a volume of 90 µL/well PBS-BN. Next, individual positive plasma samples (range of dilutions tested from 1/100 to 1/5000) and individual negative controls (at the same dilutions as the positive samples), were added per plate in a final volume of 100 µl per well. Two blank control wells with beads in PBS-BN were set up in each plate to control for background signal. Plates were incubated on a microplate shaker at 600 rpm and protected from light, and then washed three times with 200 µl/well of PBS-Tween20 0.05%, using a magnetic manual washer (Millipore, 43-285). For more accurate IgM measurements, we tested whether diluting samples 1:10 with GullSORB™ IgG Inactivation Reagent (Meridian Bioscience™) prior to testing for IgM levels could reduce high responses observed in some negative samples (27). Additionally, we tested the levels of RBD and S antibodies obtained at different plasma dilutions when incubated in a multiplex panel with additional antigen-beads including S1, S2, M and N constructs, compared to those obtained in singleplex, to check for potential interferences. Finally, since viral proteins have diverse immunogenicity, definitive plasma dilutions were established with titration experiments in individual positive and negative samples once the final multiplex antigen panel and all assay conditions had been selected.

### Secondary antibody incubation and plate reading

We compared the performance of the assays when using biotinylated secondary antibodies followed by SAPE, versus secondary antibodies conjugated directly to PE, and at different incubation times (45 versus 30 min). In all cases, each new lot of secondary antibody was titrated for selecting the optimal concentration. For the first option, 100 µL of biotinylated secondary antibody diluted in PBS-BN (anti-human IgG, B1140, 1/1250; anti-human IgM, B1265, 1/1000; or anti-human IgA, SAB3701227, 1/500; Sigma) were added to all wells and incubated for 45 min at 600 rpm at RT and protected from light. Plates were washed three times, and 100 µL of SAPE (Sigma, 42250) diluted 1:1000 in PBS-BN were added and incubated during 30 min at 600 rpm, RT and protected from light. For the second option, 100 µL of PE-secondary antibody diluted in PBS-BN (goat anti-human IgG, GTIG-001, 1/400; goat anti-human IgM, GTIM-001, 1/200; or goat anti-human IgA, GTIA-001, 1/200; Moss, MD, USA) were added to all wells and incubated for 45 or 30 min at 600 rpm at RT and protected from light.

Plates were washed three times, beads resuspended in 100 µl of PBS-BN, and data acquired using a Luminex® 100/200 analyzer with 70 µl of acquisition volume per well, DD gat 5000-25000 settings, and high PMT option. Plates could also be kept ON at 4°C, protected from light, and read the next day. At least 50 beads were acquired per antigen and sample. Crude median fluorescent intensities (MFI) were exported using the xPONENT software. Seropositivity threshold (cutoff) for optimization tests was calculated as 10 to the mean plus 3 standard deviations of log_10_-transformed MFIs of the negative controls for each antibody isotype and antigen.

### Performance of the SARS-Cov-2 qSAT assays

The Receiver operating characteristic (ROC) curves, their corresponding area under the curve (AUC), and the specificity and sensitivity of the IgM, IgA and IgG assays, were established testing all 115 positive samples from participants diagnosed with SARS-CoV-2 infection, regardless of symptoms information and at different periods since the onset of symptoms (7, 14, 21 and 28), and 128 negative samples. For IgM, IgA and IgG assays the multiplex panel including RBD, S, S1, S2, M and N antigen constructs was used, following the same procedures as indicated above and after selecting the optimal assay conditions.

## Data analysis

ROC curves and AUC, sensitivities and specificities were calculated using the predicted values estimated by supervised machine learning Random Forest (RF) algorithm models with all pre-pandemic negative controls and COVID-19 positive controls. IgM, IgA and IgG MFIs to the different antigens or their combinations were the predictors, and the outcome was SARS-CoV-2 positivity or negativity. Antibody/antigen variables that did not discriminate between positive samples from negative controls were excluded from the analysis. Then the antibody/antigen variables were further down-selected using an RF algorithm including all negative and positive controls (N=243) or all negative controls plus positive controls corresponding to each different period since onset of symptoms: ≥7 days (N=221), ≥14 days (N=207), ≥21 days (N=179), ≥28 days (N=155). The importance of the variables was ranked according to the Mean Decrease Accuracy and the Mean Decrease Gini. Next, different RFs were built exploring all possible variable combinations at the different periods based on the selected variables per period: all samples (top 12 markers), ≥7 days (top 12 markers), ≥14 days (top 10 markers), ≥21 days (top 10 markers) and ≥28 days (top 11 markers). For each model, we calculated the AUC and selected three seropositivity cutoffs aiming at specificities of i) 100%, ii) ≥ 99% and <100%, and iii) ≥ 98% and <99%, and obtained the corresponding sensitivity. Models with 100% specificity and the highest sensitivities were selected for ROC curve representations. The analysis was carried out using the statistical software R studio version R-3.5.1 (28) (packages used: randomForest (29) and pROC (30)).

## RESULTS

The characteristics of SARS-CoV-2 infected participants whose plasma samples have been used in the study, with regards to age, sex, days since rRT-PCR diagnosis and days since onset of symptoms, are included in **Table S1.**

### Selection of optimal concentration for protein coupling to microspheres

The optimal amount of protein to be coupled to beads depended on the antigen and needs to be tested with each new lot. Among the concentrations tested (10, 30 or 50 μg/mL protein), titration curves did not usually change substantially, in which case the lower concentration was chosen for the subsequent experiments. An illustrative example is shown for which the medium concentration was slightly superior when tested for IgG and IgM and thus selected **(Figure S1)**.

### Optimization of sample incubation conditions

#### Duration of incubation

To establish the optimal range of plasma dilutions for the measurement of IgM, IgA and IgG antibodies to our primary SARS-CoV-2 antigen (RBD), we initially tested positive and negative samples at four concentrations (1/100, 1/500, 1/2000, 1/5000) in singleplex **(Figure 1)**. We found 1/200 to 1/500 to be in the adequate dilution range for the subsequent assay optimization experiments. Our original standard operating procedures (SOP) established for large seroepidemiological and vaccine studies using *Plasmodium falciparum* antigens were based on ON incubations at 4°C (16). For the COVID-19 serology, we prioritized having faster assays and thus compared the performance of ON incubations at 4°C versus shorter times at RT. We tested that the range of dilutions was still adequate when reducing the incubation time **(Figure 2A)** and compared antibody levels and number of seropositive samples incubating ON at 4°C versus 2 h RT at 1/500 **(Figure 2B)**. Although the MFI readings in positive samples generally diminished with shorter times, the MFI readings in the negative samples also reduced, i.e. the signal to noise ratio was the same or sometimes better, maintaining or increasing the overall proportion of seropositive among the positive samples and thus the sensitivity. Based on these data, we adopted the 2 h incubation time for an initial COVID-19 seroprevalence study (19). We subsequently tested shorter incubations more extensively and found that 1 h was non-inferior to 2 h incubation **(Figure 2C)** and thus 1 h was selected for the optimized SOP.

**Figure 1.**
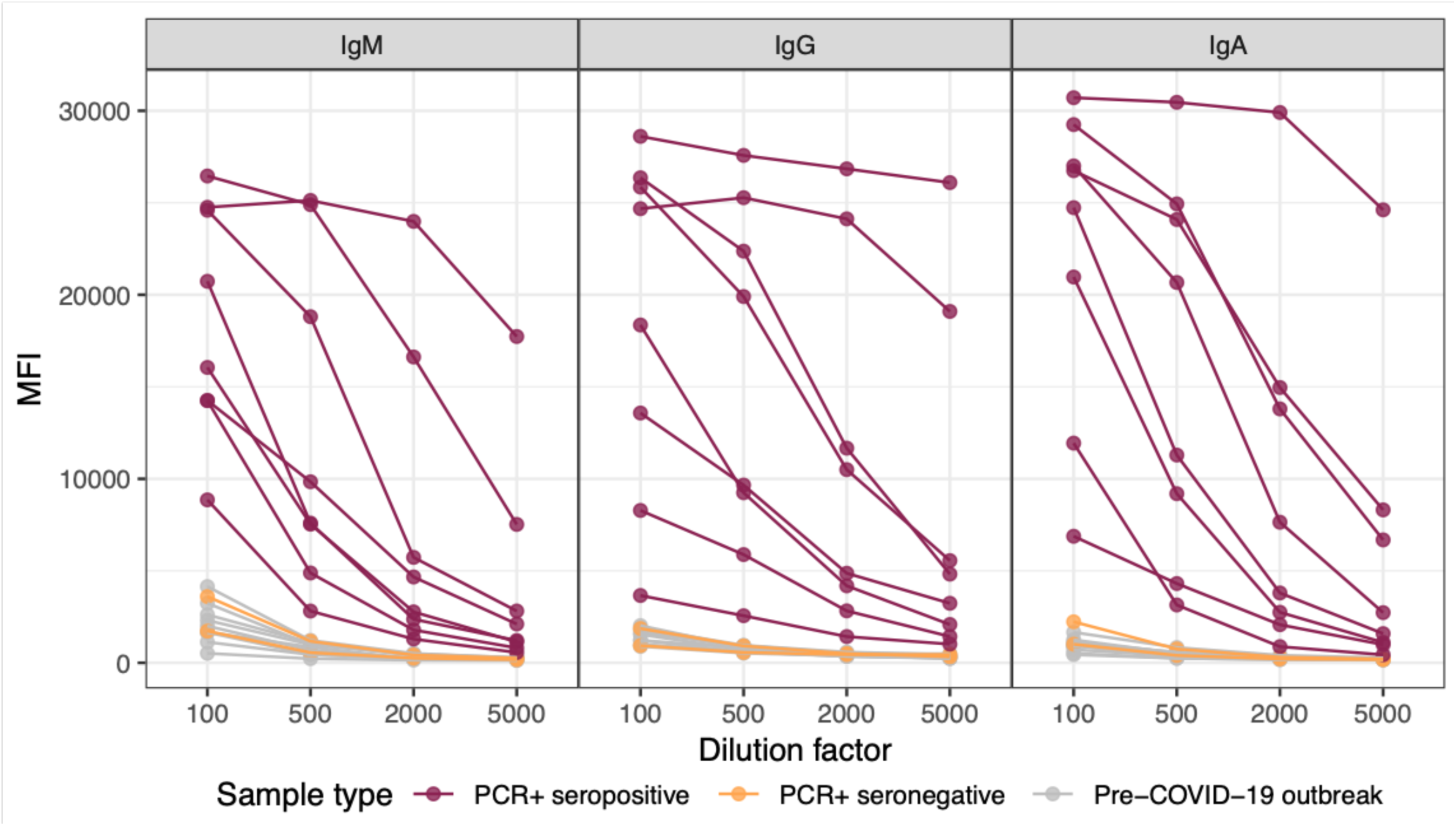
Levels (median fluorescence intensity, MFI) of IgM, IgA and IgG antibodies to RBD antigen of SARS-CoV-2 in singleplex using samples from positive and negative individuals at different dilutions after overnight incubation at 4°C.

**Figure 2.**
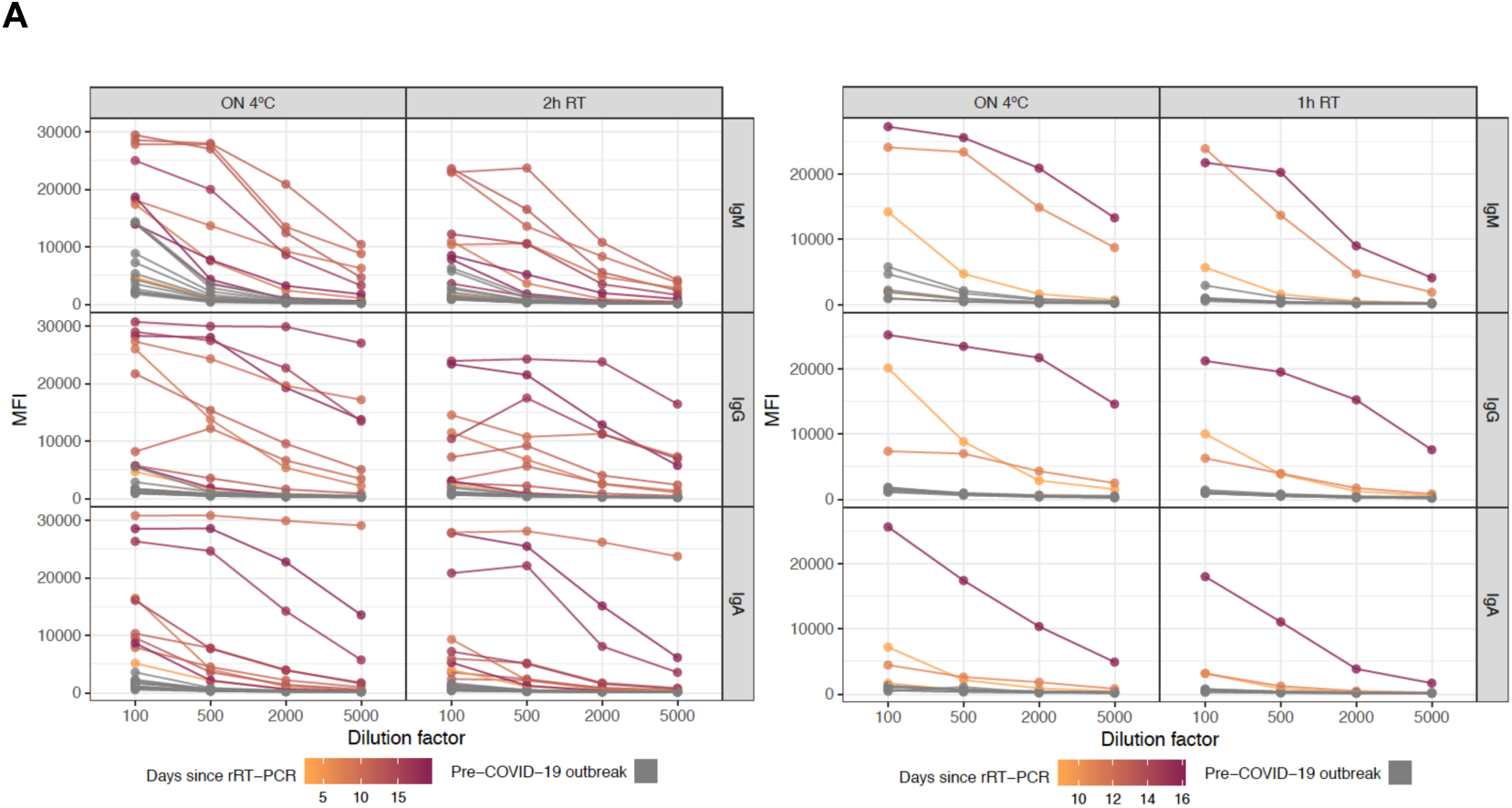
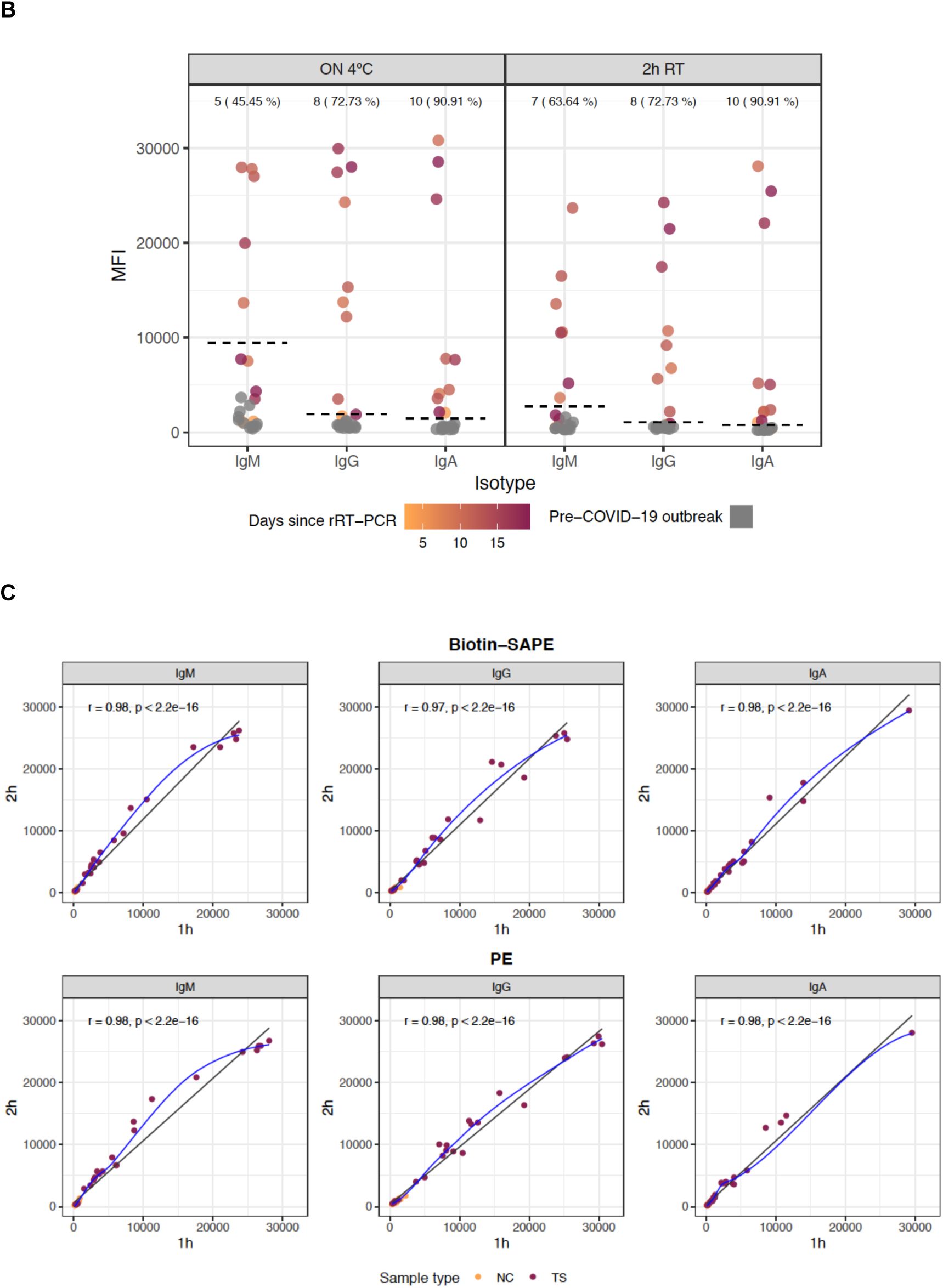
Levels of IgM, IgA and IgG antibodies (median fluorescence intensity, MFI) to RBD antigen of SARS-CoV-2 in positive and negative individuals, comparing different sample dilutions incubating overnight (ON) at 4°C versus 2 hours or 1 hour at room temperature (RT) **(A)**. One dilution (1/500) was chosen to further compare the sensitivity in the detection of positive samples with incubation ON at 4°C versus 2 hour at RT **(B)** and 2 hours versus 1 hour at RT using the two different secondary antibodies evaluated in this study **(C).** In B, Cutoff values are indicated by dashed lines. The number and percentage of seropositive samples within rRT-PCR+ is shown at the top of the dot plots. In C, the blue fitting curve was calculated using the LOESS (locally estimated scatterplot smoothing) method and the black line by linear regression. Spearman test was used to assess the correlations. NC, negative controls; TS, test samples.

#### Reduction of background in IgM assay

Treatment with GullSORB™ reduced or did not change the MFI signal, depending on the sample, antigen and dilution **(Figure S2A**). This additional incubation generally increased the signal to noise ratio and thus sensitivity and number of seropositive IgM responses among the positive controls, particularly at the lower dilutions, therefore the GullSORB™ incubation was adopted for this assay **(Figure S2B)**. IgM reactivity in negative controls was lower against S-based antigens than against M- or N-based antigens and thus GullSORB™ treatment benefited the signal to noise ratio the most in these later proteins.

#### Singleplex versus multiplex antigen testing

Multiplexing the antigens (8-plex panel) did not significantly decrease the MFI antibody levels to RBD or S compared to singleplex testing **(Figure 3A)** neither for any of the other antigens **(Figure S3A)**. Interestingly, there was no evidence of any interference between RBD, S, S1 or S2 antigens despite sharing epitopes within the same multiplex panel. A number of negative pre-pandemic samples had pre-existing antibodies recognizing SARS-CoV-2 proteins for certain isotypes and dilutions **(Figure S3A)**: IgG to S1, S2, M and N constructs, and IgA to S1 and N-term & C-term of N. Furthermore, testing plasmas against multiple antigens increased the sensitivity of the assay since some individuals who were seronegative or low responders to RBD, responded with higher antibodies to S (**Figure 3B**). Once the multiplex antigen panel was established, a set of positive and negative samples were tested at different dilution(s) covering the diverse immunogenicity of the proteins, and 1/500 and 1/3500 were selected for the assay performance evaluation **(Figures 3C and S3B)**.

**Figure 3.**
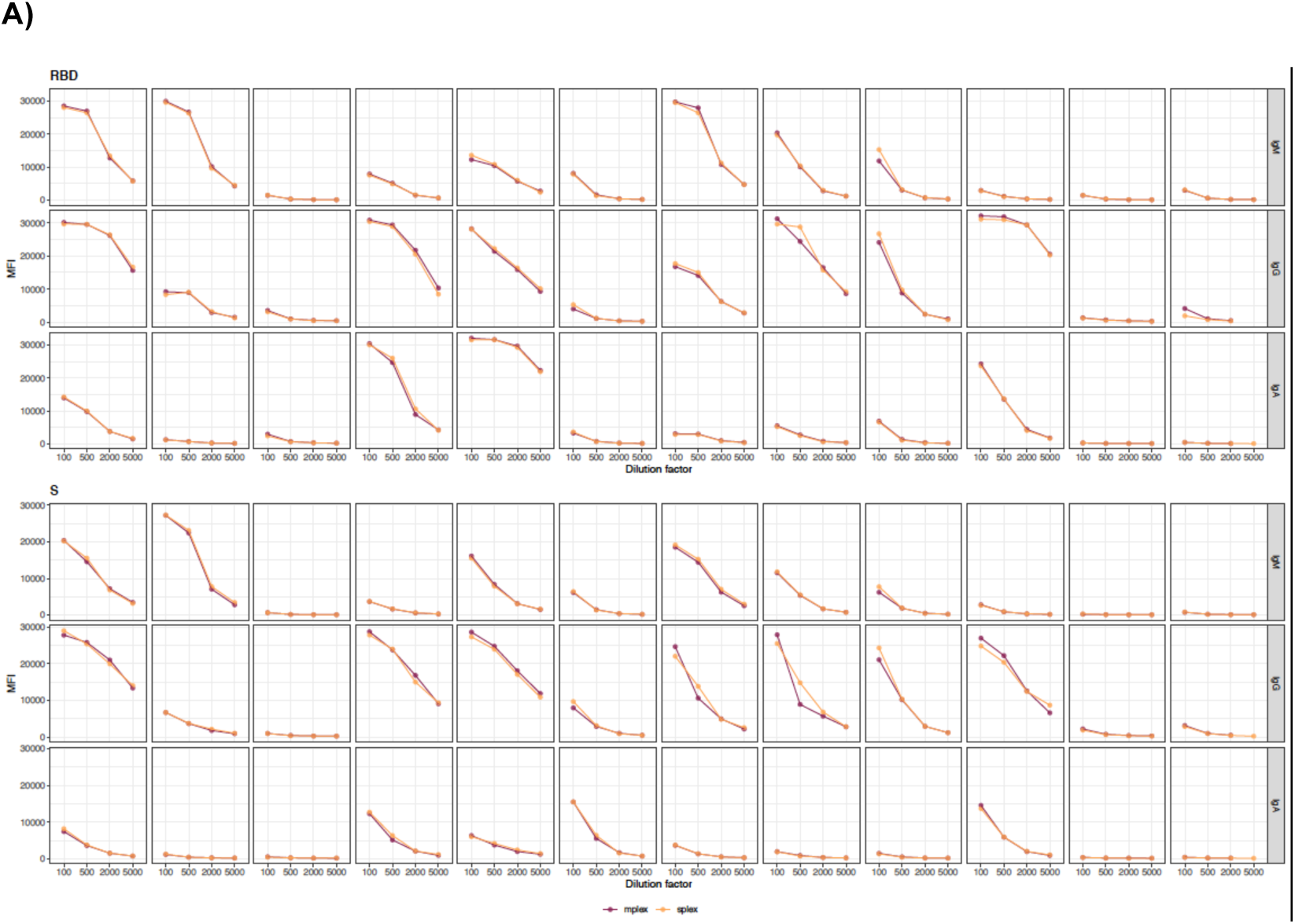
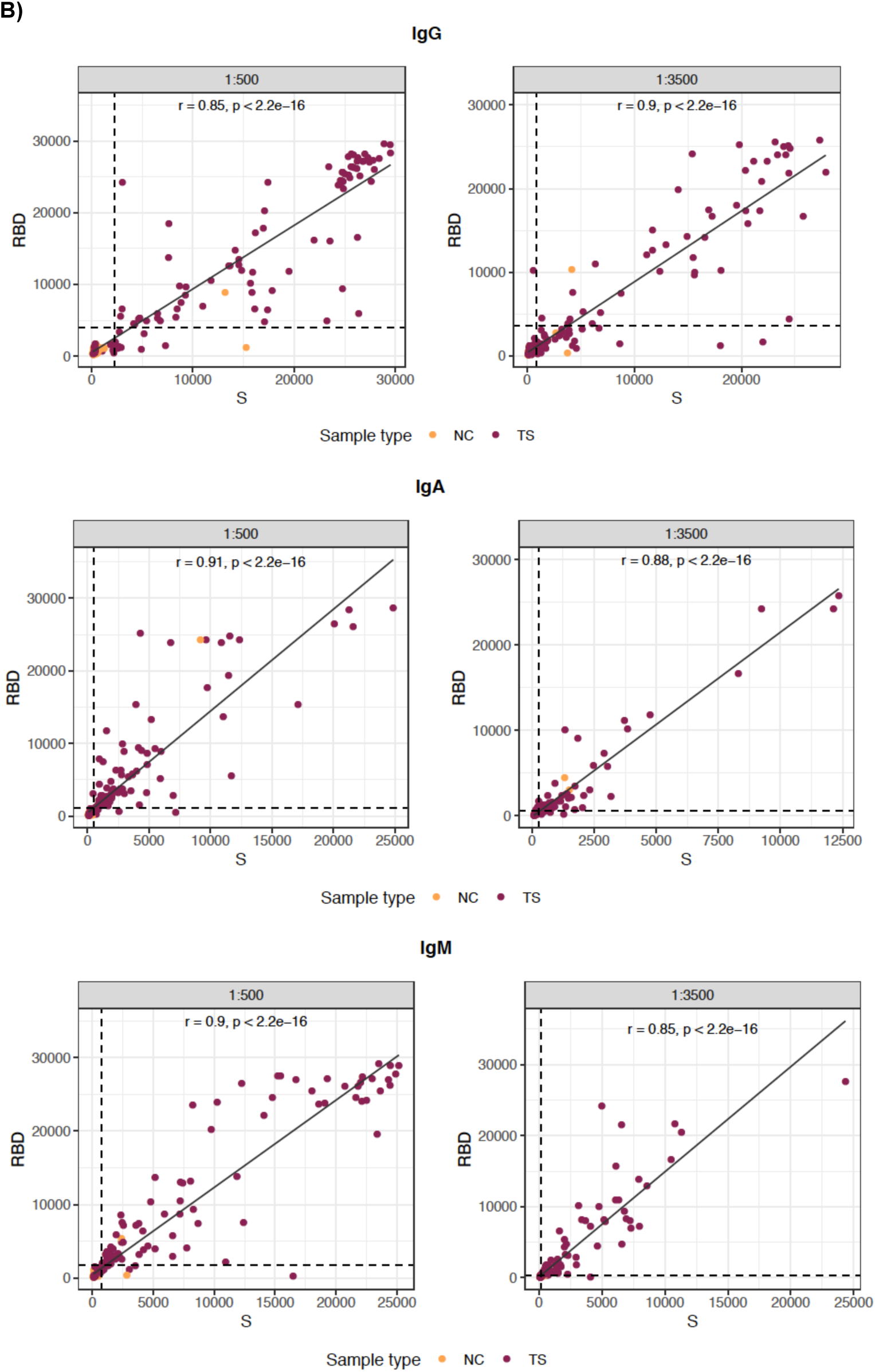
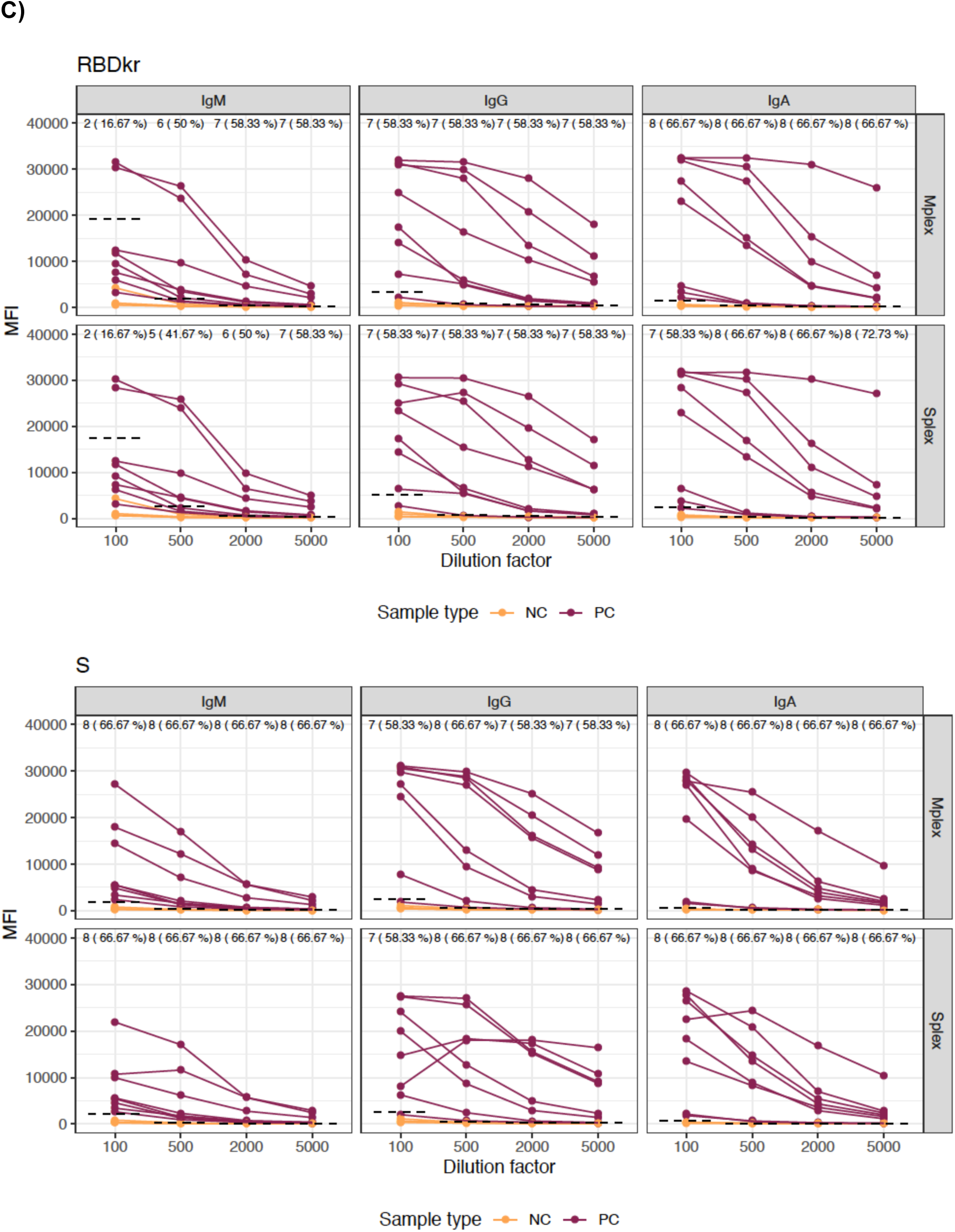
Levels of plasma IgM, IgA and IgG antibodies to the SARS-CoV-2 primary antigens spike (S) and receptor binding domain (RBD) at different dilutions. **A)** Comparison of antibody levels (MFI) in singleplex (orange) versus multiplex (burgundy); the first 10 samples from left to right are from individuals who were positive by rRT-PCR at different time periods since diagnosis, and the last two samples on the right are from individuals pre-COVID-19 pandemia (the rest of antigens are shown in Figure S3A). **B)** Correlation of IgG, IgM and IgA antibody levels against RBD versus S at different dilutions showing the benefit of including multiple antigens in the panel to maximize the detection of seropositives. **C)** Comparison of seropositivities among the positive controls tested at different dilutions in multiplex and singleplex for RBD and S antigens. Cutoff values are indicated by dashed lines. The number and percentage of seropositive samples within rRT-PCR+ is shown at the top of the dot plots. Spearman test was used to assess the correlations in C. NC, negative controls; TS, test samples.

### Optimization of secondary antibodies

Secondary antibodies conjugated to PE performed as well as a two-step secondary antibody conjugated to biotin followed by SAPE incubation **(Figure 4).** The PE-antibody reagent that resulted in a shorter assay was selected as the preferred option. Finally, 30 min incubation was non-inferior to 45 min incubation **(Figure S4).**

**Figure 4.**
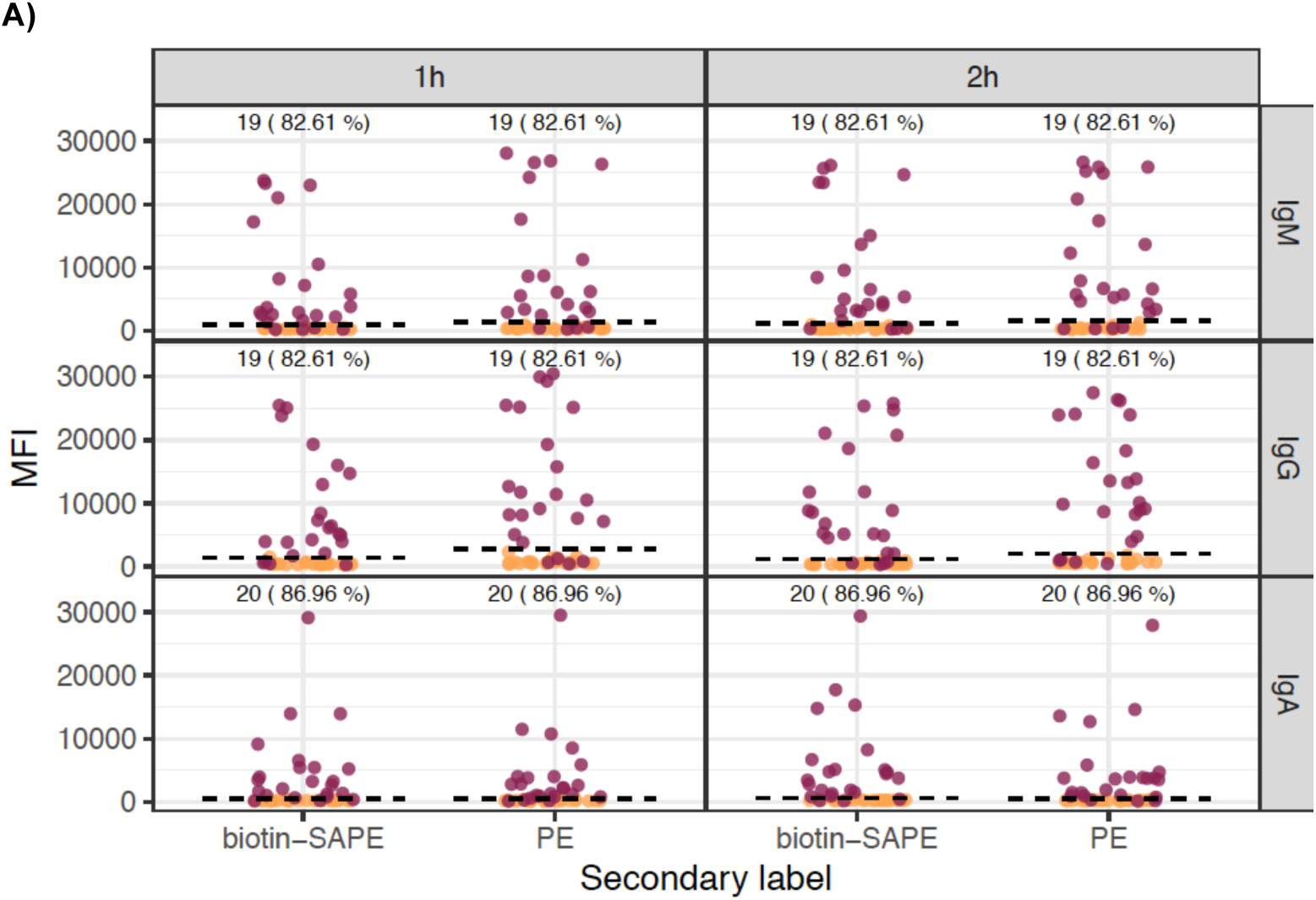
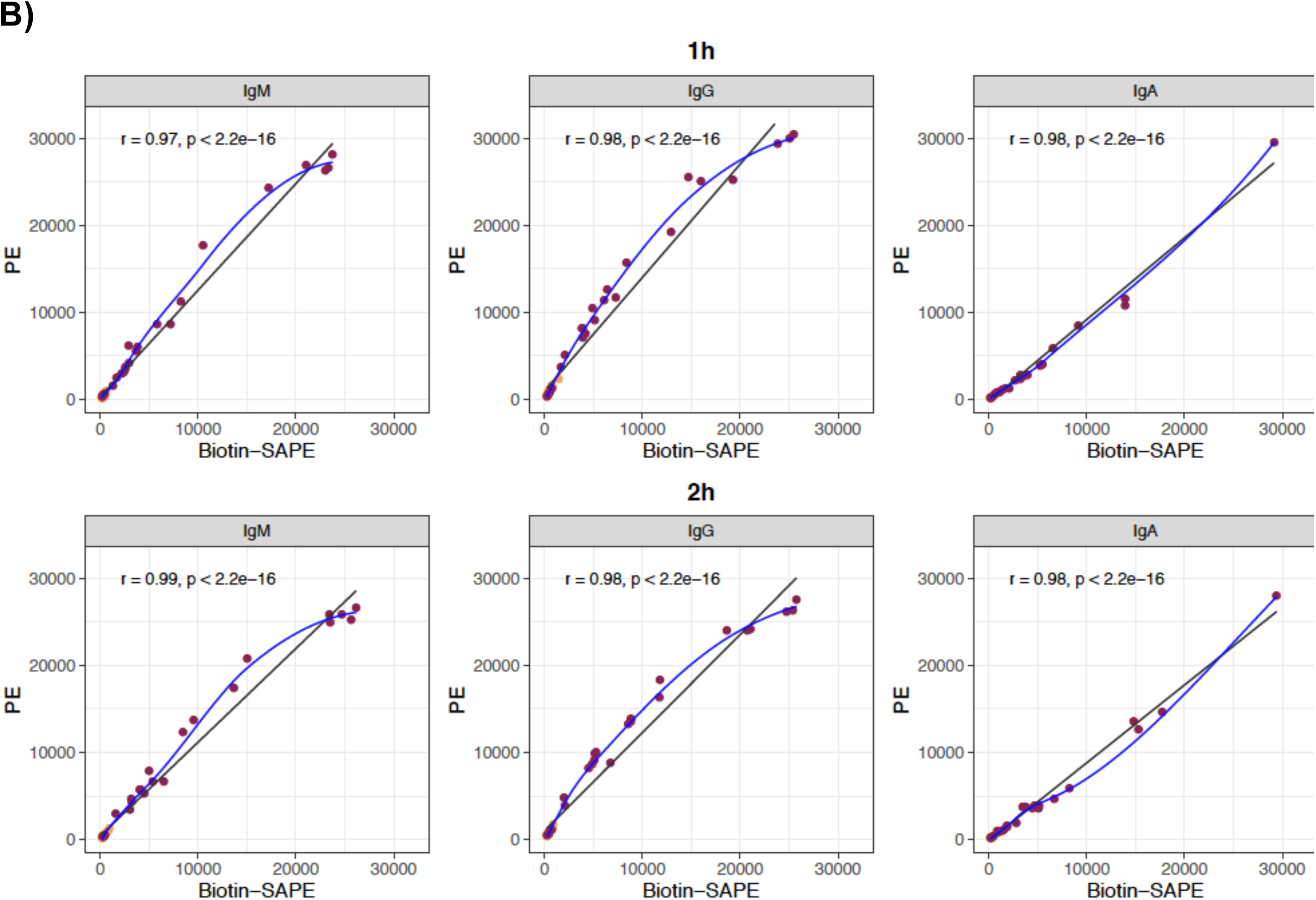
**A)** Levels of IgM, IgA and IgG antibodies (median fluorescence intensity, MFI) and % seropositivity to RBD among positive controls (burgundy), comparing secondary antibodies conjugated to biotin and streptavidin-phycoerythrin (SAPE) versus PE. Negative controls in orange. **B)** Correlations between antibody levels measured using secondary antibodies conjugated to biotin and SAPE versus PE, for 1 h and 2 h sample incubations. Seropositivity cutoff values are indicated by dashed lines. The number and percentage of seropositive samples among rRT-PCR+ is shown at the top of the dot plots. In B, the blue fitting curve was calculated using the LOESS (locally estimated scatterplot smoothing) method and the black line by linear regression. Spearman test was used to assess the correlations.

### Sensitivity and specificity of the qSAT assays

We sought for the combination of Ig and antigen responses that yielded the highest specificity (primarily), sensitivity and AUC to detect seropositive responses. For RBD and S, IgG and IgA at 1/500 dilution, and IgM responses at 1/3500, gave higher percentages of seropositive responses among the positive controls and thus were selected for the calculations; for N constructs, IgG and IgA performed better at 1/3500 except for N C-term in which IgG was better at 1/500. Antibodies to M, S1 (IgG & IgA) and N N-term (IgM) did not discern well positive from negative responses and were not included in the RF models. The contribution of each antibody/antigen variable was ranked according to an RF algorithm at different periods since onset of symptoms (**Figure S5**) and the top 10-12 variables were selected. We performed RF for all the combination of variables and assessed the sensitivity of each combination at three different seropositivity thresholds aiming at specificities of 100%, 99% and 98%. The specificity of the qSAT assays in samples from participants with SARS-CoV-2 positive diagnosis with ≥14 days since the onset of symptoms (n=207) was up to 100% with sensitivity up to 94.94%, and AUC up to 0.992, for the best combinations of Ig isotypes/antigens. The top 5 performing antibody signatures for three different seropositivity thresholds targeting specificities of 100%, 99% and 98% are shown in **Table 1**, and their ROC are shown in **Figure 5**. In samples from participants with ≥21 days since the onset of symptoms (n=179), the specificity was up to 100% and the sensitivity up to 96.08%, with AUC up to 0.999 for the best combinations of Ig isotypes/antigens (**Table 2, Figure 5**). In samples from all participants regardless of time since symptoms onset (n=243), the specificity was up to 100% and the sensitivity up to 82.61%, depending on the combinations of Ig isotypes/antigens, with AUC up to 0.918 for the best combinations (**Table S2, Figure 5**). The performance of the qSAT assays to predict positivity was clearly superior using combinations of multiple Ig isotypes/antigens to using single isotype/antigen markers (**Figure 5**). Higher sensitivities were obtained when specificities were set to 98% or 99% (**Tables 1, 2 & S2**), reaching 100% for samples ≥21 or ≥28 days since the onset of symptoms.

**Table 1.** **Sensitivity and specificity of the Luminex antibody assays** at ≥14 days since onset symptoms at different thresholds targeting specificities of 100%, 99% and 98%. The top 5 performing signatures per each category are shown.

| Antibody/antigen combinations | AUC | Specificity | Sensitivity |
| --- | --- | --- | --- |
| IgA S2 + IgG N + IgG S + IgM RBD + IgM S + IgM S2 | 0.992 | 100% | 94.94% |
| IgA S + IgG N + IgG S + IgM RBD + IgM S2 | 0.991 | 100% | 94.94% |
| IgG N + IgG S + IgM RBD + IgM S + IgM S2 | 0.991 | 100% | 94.94% |
| IgG S + IgM RBD | 0.990 | 100% | 94.94% |
| IgA S2 + IgG N + IgG N Ct + IgG S + IgM RBD + IgM S2 | 0.990 | 100% | 94.94% |
| IgG RBD + IgG S | 0.984 | 99.22% | 96.20% |
| IgG N + IgG S2 + IgM S | 0.979 | 99.22% | 96.20% |
| IgG N + IgG S + IgG S2 | 0.979 | 99.22% | 96.20% |
| IgG N + IgG N Ct + IgG S + IgG S2 | 0.978 | 99.22% | 96.20% |
| IgG N + IgG S | 0.978 | 99.22% | 96.20% |
| IgG N + IgG N Ct + IgG RBD + IgG S + IgG S2 + IgM S | 0.990 | 98.44% | 97.47% |
| IgG N + IgG RBD + IgG S + IgM S | 0.989 | 98.44% | 97.47% |
| IgA S + IgG N + IgG S + IgG S2 + IgM S | 0.988 | 98.44% | 97.47% |
| IgG N + IgG S + IgG S2 + IgM S | 0.988 | 98.44% | 97.47% |
| IgG N + IgG N Ct + IgG S + IgG S2 + IgM S | 0.988 | 98.44% | 97.47% |

**Table 2.** **Sensitivity and specificity of the Luminex antibody assays** at ≥21 days since onset symptoms at different thresholds targeting specificities of 100%, 99% and 98%. The top 5 performing signatures per each category are shown.

| Antibody/antigen combinations | AUC | Specificity | Sensitivity |
| --- | --- | --- | --- |
| IgG N + IgG S + IgM RBD + IgM S2 | 0.999 | 100% | 96.08% |
| IgG N + IgG S + IgM S + IgM S2 | 0.999 | 100% | 96.08% |
| IgA RBD + IgG N + IgG S + IgM RBD + IgM S2 | 0.999 | 100% | 96.08% |
| IgG N + IgG S + IgM RBD | 0.999 | 100% | 96.08% |
| IgA RBD + IgG N + IgG N Ct + IgG S + IgM RBD + IgM S + IgM S2 | 0.998 | 100% | 96.08% |
| IgG N + IgG S2 + IgM S | 0.988 | 99.22% | 98.04% |
| IgG N + IgG S2 | 0.984 | 99.22% | 98.04% |
| IgG N + IgG S + IgM RBD + IgM S2 | 0.999 | 99.22% | 96.08% |
| IgG N + IgG S + IgM RBD + IgM S | 0.999 | 99.22% | 96.08% |
| IgG N + IgG S + IgM S + IgM S2 | 0.999 | 99.22% | 96.08% |
| IgG N + IgG S + IgM RBD + IgM S2 | 0.999 | 98.44% | 100% |
| IgG S + IgG S2 + IgM RBD + IgM S | 0.999 | 98.44% | 100% |
| IgA RBD + IgG RBD + IgG S + IgM S + IgM S2 | 0.999 | 98.44% | 100% |
| IgG N + IgG S + IgG S2 + IgM RBD + IgM S2 | 0.999 | 98.44% | 100% |
| IgG N + IgG S + IgG S2 + IgM RBD + IgM S + IgM S2 | 0.999 | 98.44% | 100% |

**Figure 5.**
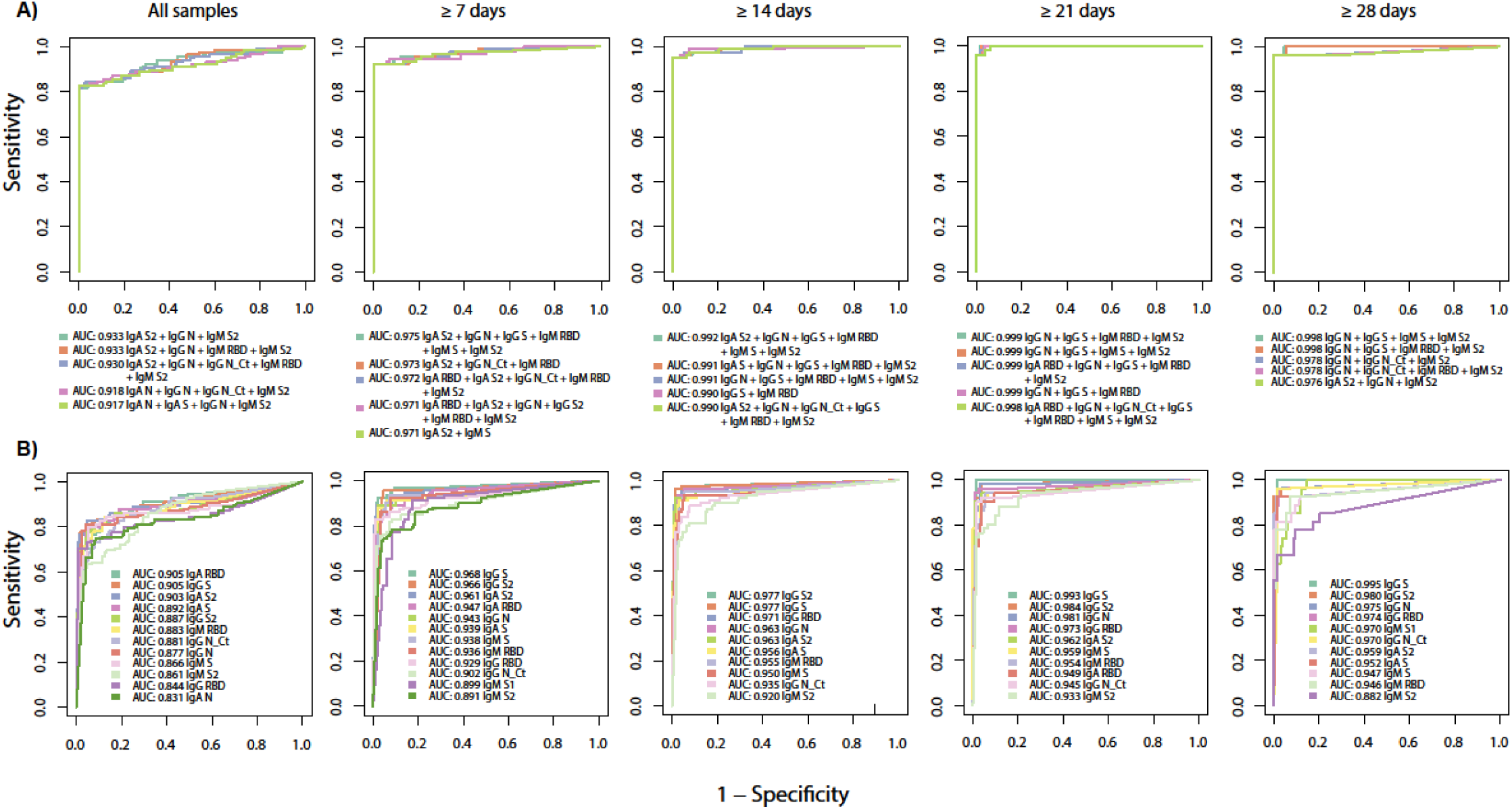
Antibody Luminex assays performance. Receiver operating characteristic (ROC) curve and area under the curve (AUC) using samples from pre-pandemic negative controls plus either all participants with positive COVID-19 diagnosis or participants with positive diagnosis at different times since onset of symptoms, comparing combinations of multiple immunoglobulin isotypes to different antigens with top performances **(A)** versus those of single isotype/antibody markers **(B).**

## DISCUSSION

We developed three novel multiplex immunoassays for quantifying IgM, IgA and IgG to eight SARS-CoV-2 protein constructs and evaluated by machine learning classification algorithms the performance of several isotype/antigen combinations to detect any positive antibody response to infection, obtaining specificities of 100% and sensitivities of 94.94% (≥14 days since symptoms onset) or 96.08% (≥21 days since symptoms onset), and very high predictability (AUC ≥0.99). Our qSAT assays, based on the xMAP technology, provide the best precision, accuracy and widest range of detection compared to classical qualitative (RDT) or quantitative (ELISA) assays.

For any given test, there is usually a trade-off between sensitivity and specificity. To evaluate the performance of the assays here, we prioritized specificity over sensitivity for the implications that false positives may pose at a personal level and the impact that specificity has in seroprevalence studies. Particularly when prevalence of infection is low, the positive predictive value of a test strongly relies on a high specificity. For example, in a scenario of 5% prevalence and 95% sensitivity, the positive predictive value of the test decreases from 100% to 50%, with a reduction in specificity from 100% to 95%. However, other seropositivity thresholds can be used to have a balanced specificity/sensitivity or to maximise sensitivity.

A time period after the onset of symptoms is usually established for these analyses, because antibodies take an average 4-14 days since infection to be produced and detected depending on the isotype and test (IgM 5-12 days, IgA 5-12 days, IgG 4-14 days) (31)(32)(33)(34)(35)(36). Thus, it is not necessarily expected to detect antibodies in individuals who are acutely infected and diagnosed around the time of plasma collection. Accordingly, when considering all samples, which included 13 and 14 individuals with less than 6 and 14 days since onset of symptoms, respectively, sensitivity was lower (up to 82.61%) at specificities of 100%. However, we detected IgM or IgA as early as 2 days, and IgG as early as 1 day, from onset of symptoms. In fact, since samples were collected in the early days of the COVID-19 pandemic, it is expected that IgM and IgA, which are induced upon primary infection earlier than IgG, could contribute to a higher sensitivity of detection. Most of the best signatures identified included IgM and IgA besides IgG, regardless of the time period since onset of symptoms, also beyond 28 days. However, over time, the only antibodies that would be expected to remain in blood are IgG due to the decay of IgM and IgA, e.g. IgM levels may become undetectable by the fifth week after symptoms onset (37). Therefore, with longer days since infection, the serological assays to detect maintenance of antibodies could focus on IgG detection.

The superior performance of the qSAT assays is partly based on direct fluorescence detection as opposed to colorimetric detection mediated by an enzyme. Also, antigens are covalently coupled to beads as opposed to passive coating of the ELISA plates, leading to a higher density of antigen per surface area and less antigen wash off during the assay. The higher background of ELISA microplates is related to the fact that they have a much larger surface area than the combined area of 2500 microspheres, which is more prone to the binding of non-specific antibodies if blocking is not performed correctly (14). The sensitivities and specificities of other SARS-CoV-2 serological assays externally validated with >100 positive and >100 negative samples (as recommended by FIND protocols), some of them approved by the USA FDA, are summarized in **Table S3** (38)(39).

While Luminex assays generally have high correlation to ELISAs in singleplex (R^2^ ∼0.9) (13), it is important that the assays perform equally well in multiplex format, with no interference noted between antigens, even if they had overlapping epitopes. A key value of multiplexing is that it allows to capture a wider breadth of responses and this is needed because some individuals may not respond to one antigen (e.g. RBD) but may do so to other antigens (e.g. S or N proteins) (40)(41)(42). Here, we substantially increased the sensitivity of the assay when combining isotypes/antigens compared to using only one isotype/antigen. The addition of N was more beneficial to detect seropositive responses when the onset of symptoms was recent, as this antigen is the most abundant and immunogenic and specific antibodies appear to be elicited earlier (43). In contrast, combinations of S antigens seemed to be sufficient to detect seropositive responses with longer periods since the onset of symptoms. An added advantage of multiplexing is the reduced usage of sample volume, resources and time, if antibodies to several antigens are to be evaluated. The possibility to perform miniaturized assays in small amounts of blood is very attractive in paediatric studies, in large field surveys where fingerpick may be more logistically feasible, and to test special tissues of interest including mucosal fluids.

Those combined advantages have a direct impact on the cost-efficacy of the qSAT assay, that is overall cheaper than RDT or ELISA assays. The cost of the xMAP assay can be less than one-fifth of the least expensive commercial ELISA and less than one-sixteenth of the most expensive commercial kit. Cost is reduced because there is less protein used due to the smaller surface area and less amounts of other materials and reagents. We reduced the dilutions of plasma and titrated the secondary antibody to use the minimal amounts of samples and reagents, without compromising sensitivity. The economy of scale will improve further when the assays are adapted to high throughput FlexMap3D 384-well plate format but they are also easily adaptable to the bench top MagPix 96-well format that is more affordable and easy to maintain even in remote laboratory settings.

Interestingly, positive antibody responses to M, S1, S2 and N antigen constructs were detected in samples collected before the COVID-19 pandemic. The presence of such antibodies has been interpreted as cross-reactivity with antigens of coronaviruses causing the common cold (10)(44)(45). Indeed, higher sequence homology at the protein level between SARS-CoV-2 and coronaviruses has been reported for N (particularly N-terminal and central regions), M and S2 (10)(46)(47). Pre-existing SARS-CoV-2-specific T cells have been recently reported and also attributed to cross-reactivity to human coronaviruses previously encountered (48)(49). The multiplex nature of the assay will allow to test this hypothesis in the future with the addition of antigens to related coronaviruses 229E, HKU1, NL63 and OC43 in the same assay panel, by comparing the patterns of antibody reactivity, in order to address the significance of this in immunity to COVID-19.

Here, antibody responses to M were very marginal and did not contribute to higher assay sensitivity and this could partly be because the purity of the protein was not high. However, this antigen may be valuable in studies establishing the antibody correlates of protection since at present the targets of immunity have not been elucidated. It is possible that, in addition to neutralizing antibodies directed to the RBD region of S, antibodies of other specificities with non-neutralizing functions, for example Fc-mediated opsonisation and phagocytosis, could be relevant in protection. In fact, T cell responses to epitopes located on M have been detected at high frequencies (48), and it is possible that antibodies to this or other less immunogenic antigens may also have a role in protection in some individuals. In our study, the addition of S1 from a commercial supplier did not have any added value but for future versions of the assay we will test S1 from different sources, as this subunit is expected to not cross-react with other beta-coronaviruses and be specific for SARS-CoV-2 diagnostics (11)(46).

The assays performances were excellent but further testing needs to be performed with longer periods of time since onset of symptoms, although we do expect to maintain high specificity and sensitivity albeit antibody signatures would be different and based on IgG only. Future studies will include additional positive samples of asymptomatic individuals, who probably have lower antibody levels than mild or severe cases and are rarely included in the validation of commercial kits. In addition, it will be interesting to include negative controls reacting with other coronaviruses or other infections (e.g. malaria) and pathologies known to induce polyclonal responses or rheumatoid factor, which may increase background responses.

In conclusion, we developed 100% specific and fast assays with possibly one of the best diagnostic characteristics reported in the published literature to assess seroprevalence of COVID-19. Considering their high sensitivity, these qSAT assays would be suited to identify individuals with levels of antibodies below the lower limit of detection of RDT or the lower limit of quantification of ELISA, such as asymptomatic children or immunosuppressed individuals, or long-term decaying antibodies (50). In addition this approach would be particularly suited to identify hyper immune donors with very high levels of antibodies and the largest antigenic breadth for immunotherapy. The assays are highly versatile, being easily adaptable to quantify other antibody IgG and IgA subclasses and avidity with the use of chaotropic agents, and even functional activity like binding inhibition to the virus receptor ACE2. The multiplex capabilities make them also ideal for sizeable peptide screenings to accelerate epitope mapping and selection for identifying fine-specificity of immune correlates of protection for vaccine development, and would also be applicable in vaccine evaluation when the first candidates reach larger-scale phase 2 and 3 clinical trials.

## Acknowledgements

We thank the volunteers who donated blood for COVID-19 studies and the clinical and laboratory staff who participated in the sample collection and processing. Special thanks to ISGlobal colleagues P. Cisteró, R.A. Mitchell, C. Jairoce, S. Alonso, J. Moreno, L. Puyol, C. Chaccour, and those involved in data analysis and/or recruitment of volunteers at the hospital, J.L. del Pozo, M. Fernández, M. Tortajada, C. Guinovart, S. Sanz, S. Méndez, A. Llupià, E. Chóliz, A. Cruz, S. Folchs, M. Lamoglia, N. Ortega, N. Pey, M. Ribes, N. Rosell, P. Sotomayor, S. Torres, S. Williams, S. Barroso, A. Trilla and P. Varela. We are grateful to F. Krammer for donation of RBD and S plasmids, to L. Mayer for assistance with literature review, and to Wilco de Jager from Luminex for technical advice.

