## Supplementary material for "Highly sensitive and specific multiplex antibody assays to quantify immunoglobulins M, A and G against SARS-CoV-2 antigens"

#### SUPPLEMENTARY FIGURES

**Figure S1.** Selection of the optimal RBD antigen coupling concentration. **A)** Comparison of titration curves with an anti-histidine tag biotinylated antibody at different protein concentrations (left panel) and with a positive plasma pool for IgG and IgM (right panel). **B)** Comparison of IgG and IgM titration curves with a positive plasma pool against 5 different batches of 30 μg/mL RBD-coupling reactions (A-E) for IgM **(C)** and IgG **(D)** showing highly reproducible titration curves.

**A)**

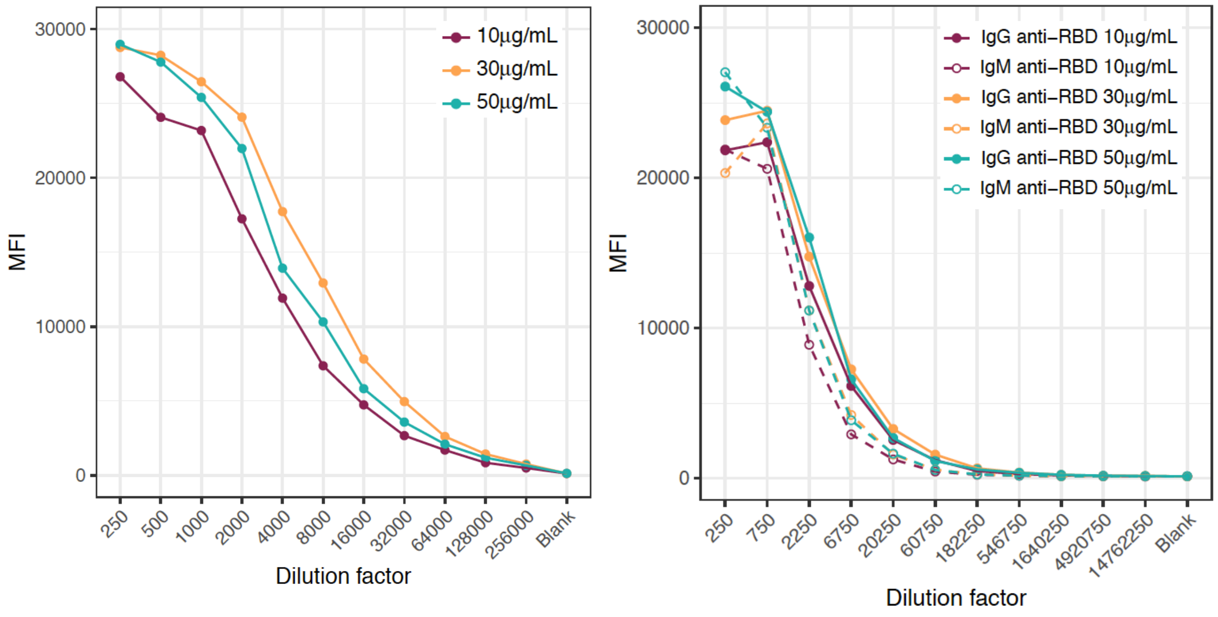

**B)** **IgM** **IgG**

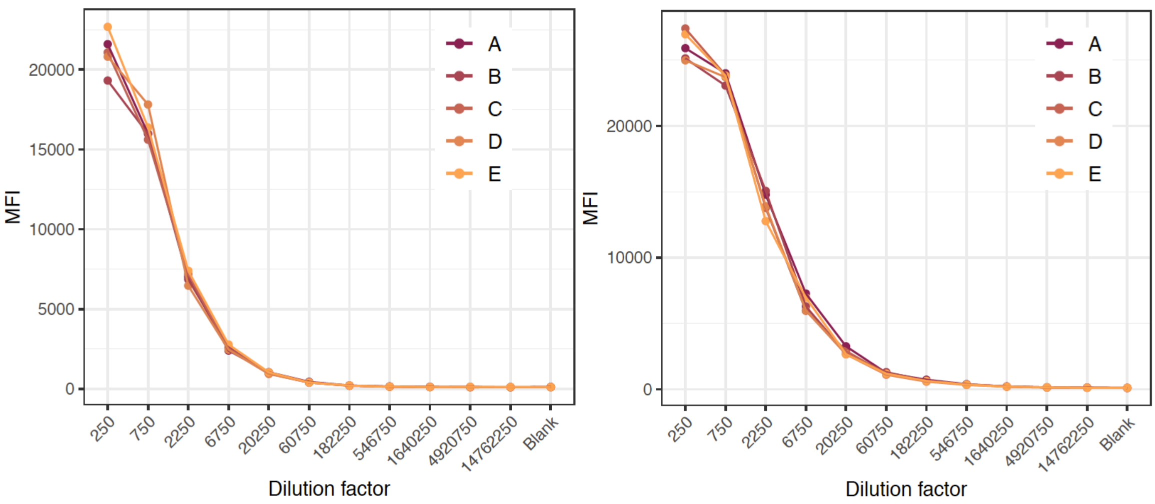

**Figure S2.** Effect of GullSORB™ treatment on IgM antibody levels to negative (NC) and positive (PC) control samples in a multiplex antigen panel at different plasma dilutions.

**A)**

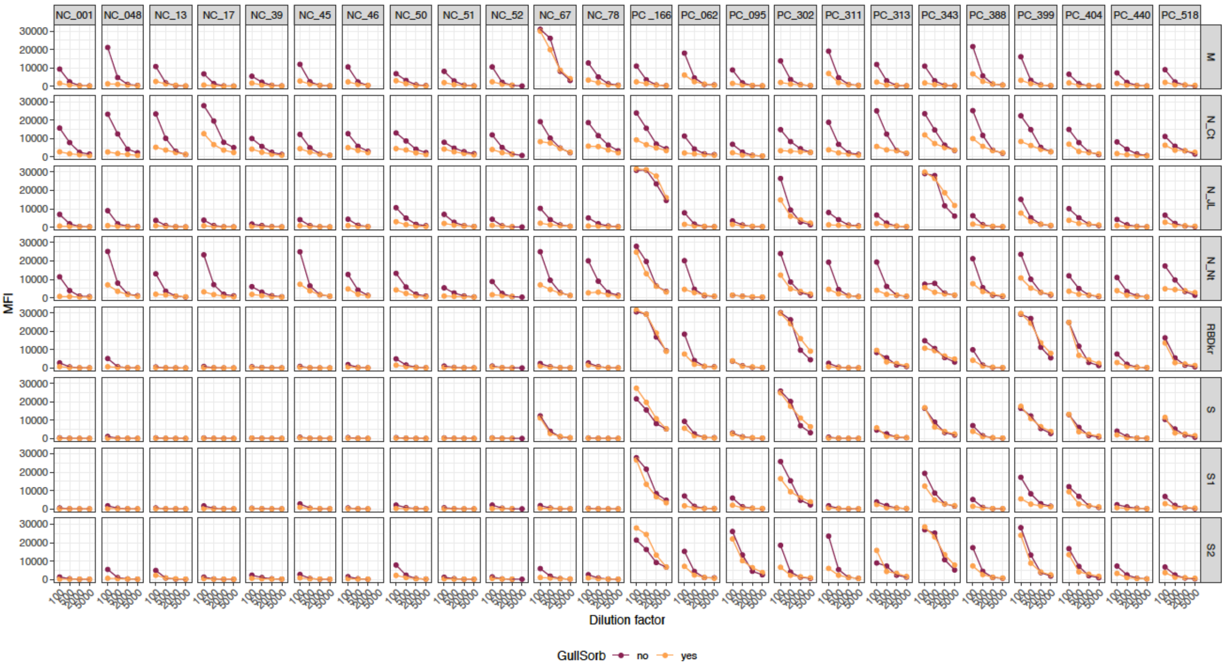

**B)**

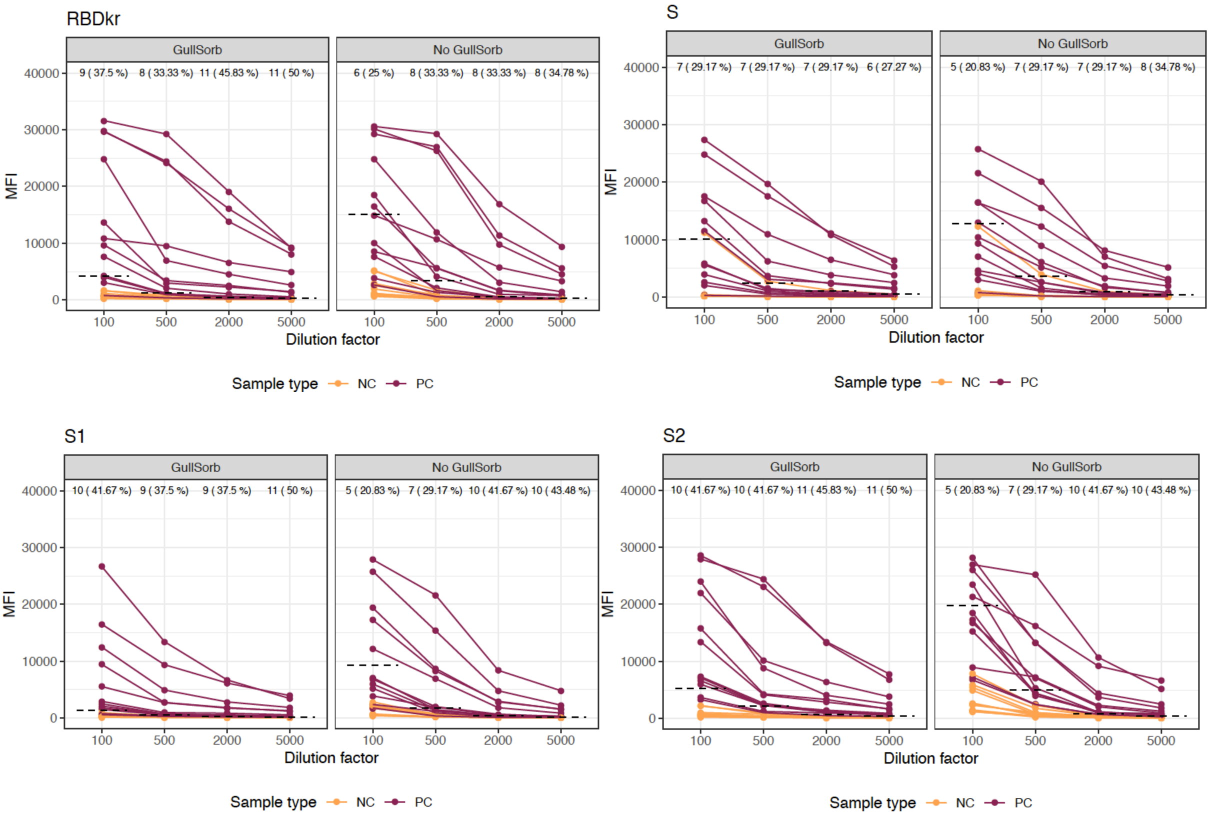

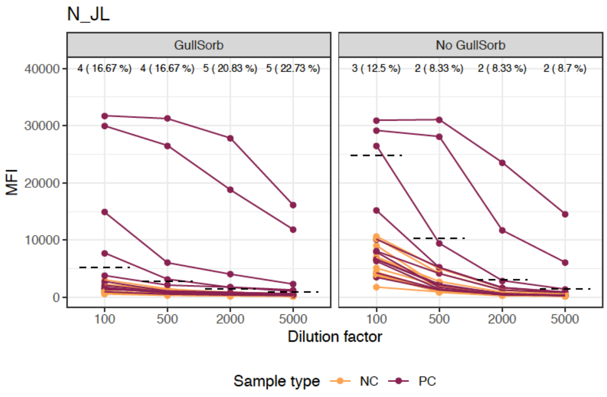
N_JL
GullSorb
No GullSorb
4 ( 16.67 %)
4 ( 16.67 %)
5 ( 20.83 %)
5 ( 22.73 %)
3 ( 12.5 %)
2 ( 8.33 %)
2 ( 8.33 %)
2 ( 8.7 %)
40000
30000
20000
10000
0
MFI
100
500
2000
5000
Dilution factor
Sample type
NC
PC

**Figure S3. A)** Comparison of IgM (without GullSORB), IgA and IgG antibody levels (MFI) in plasma samples with antigen coated microspheres in singleplex (yellow) versus multiplex (red). The first 4 samples from left to right correspond to negative individuals and the remaining 8 samples to positive individuals. **B)** Comparison of seropositivities for each isotype at different dilutions of positive (red) or negative (yellow) plasmas for antigen N.

**A)**

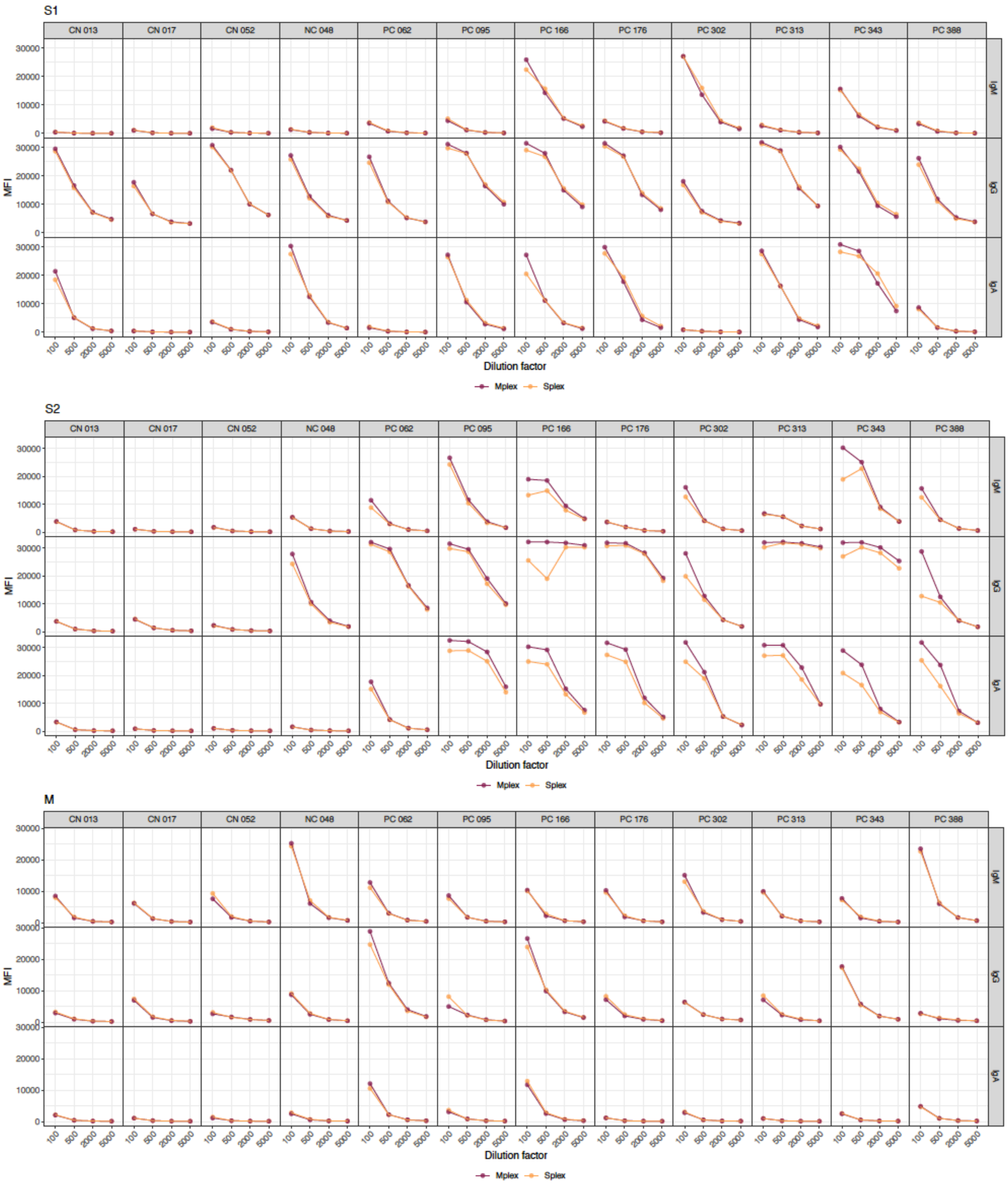

N_JL

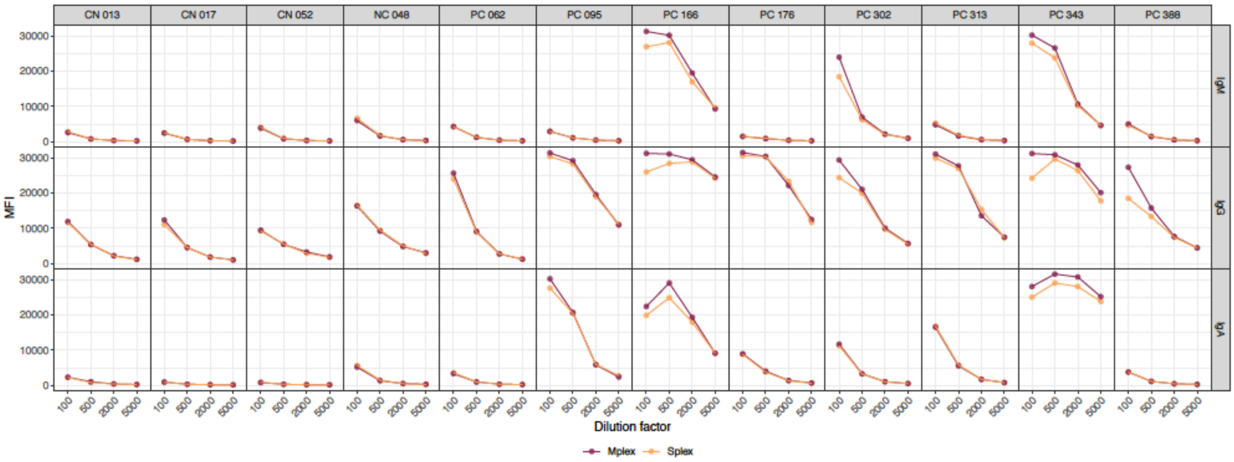
CN 013
CN 017
CN 052
NC 048
PC 062
PC 095
PC 166
PC 176
PC 302
PC 313
PC 343
PC 388
IgM
IgG
IgA
MFI
Dilution factor
Mplex
Splex

N_Ct

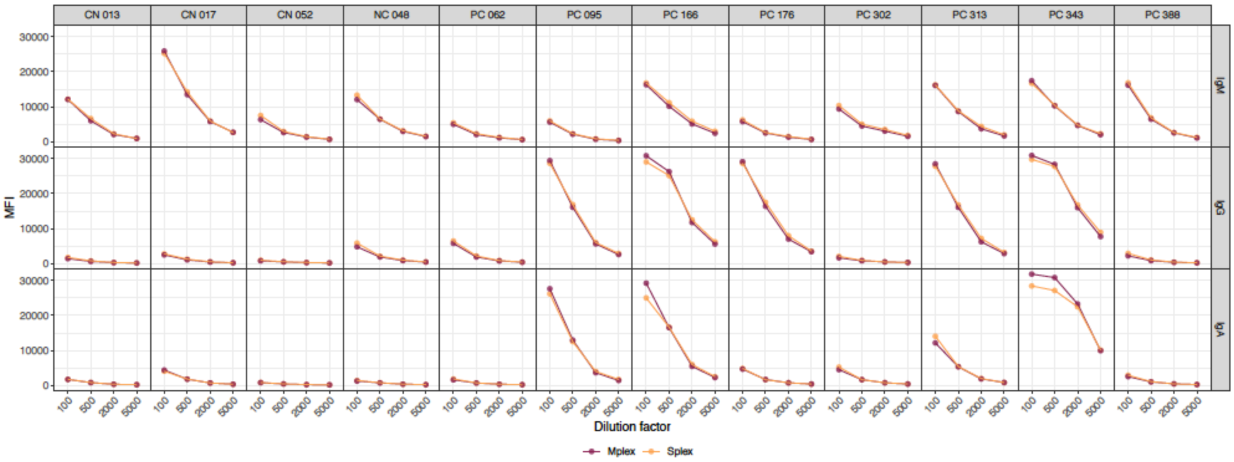
CN 013
CN 017
CN 052
NC 048
PC 062
PC 095
PC 166
PC 176
PC 302
PC 313
PC 343
PC 388
IgM
IgG
IgA
MFI
Dilution factor
Mplex
Splex

N_Nt

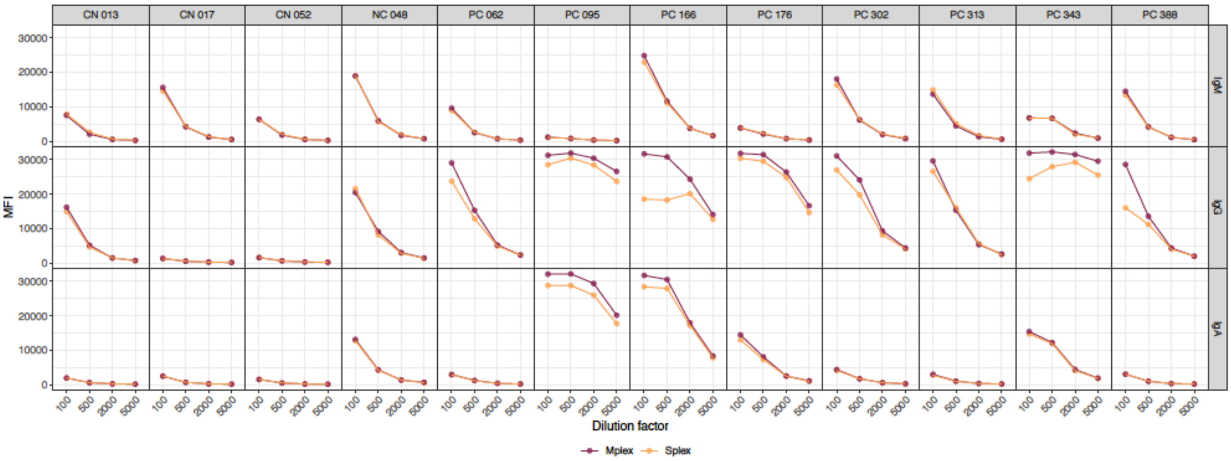
CN 013
CN 017
CN 052
NC 048
PC 062
PC 095
PC 166
PC 176
PC 302
PC 313
PC 343
PC 388
IgM
IgG
IgA
MFI
Dilution factor
Mplex
Splex

**B)**

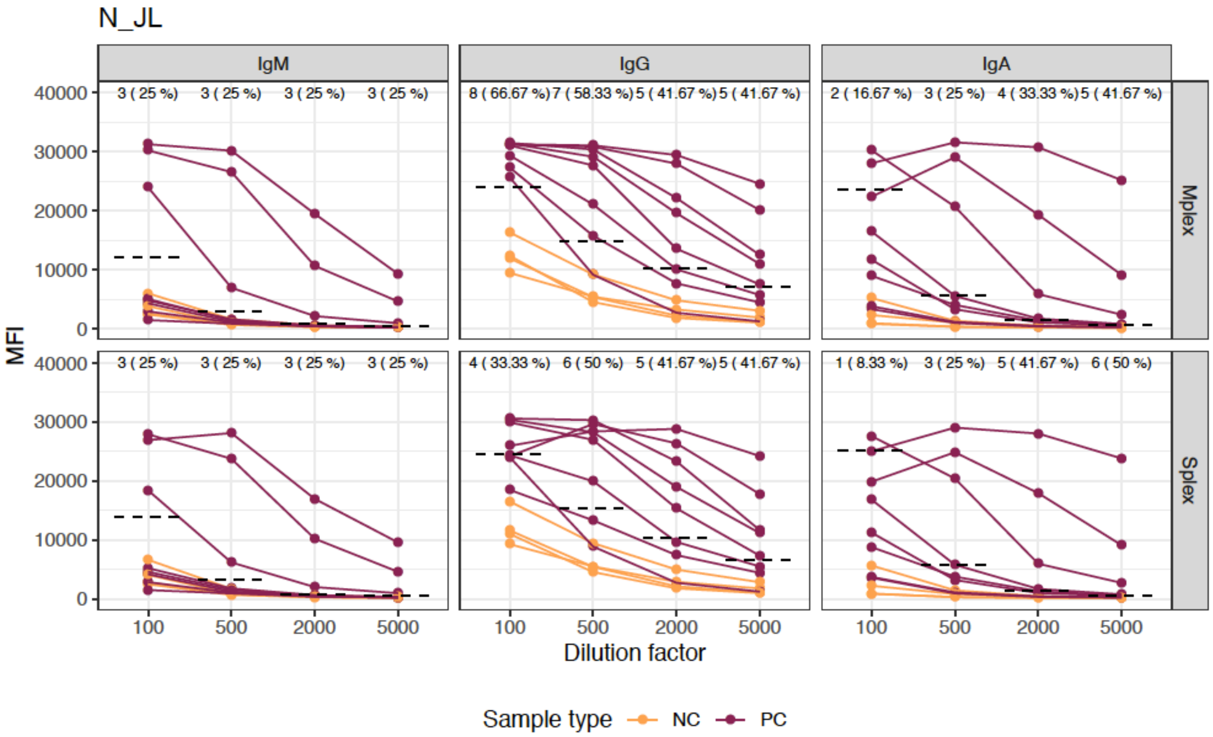

N_JL
IgM
IgG
IgA
Mplex
Splex
3 ( 25 %) 3 ( 25 %) 3 ( 25 %) 3 ( 25 %)
8 ( 66.67 %) 7 ( 58.33 %) 5 ( 41.67 %) 5 ( 41.67 %)
2 ( 16.67 %) 3 ( 25 %) 4 ( 33.33 %) 5 ( 41.67 %)
3 ( 25 %) 3 ( 25 %) 3 ( 25 %) 3 ( 25 %)
4 ( 33.33 %) 6 ( 50 %) 5 ( 41.67 %) 5 ( 41.67 %)
1 ( 8.33 %) 3 ( 25 %) 5 ( 41.67 %) 6 ( 50 %)
MFI
40000
30000
20000
10000
0
100
500
2000
5000
Dilution factor
Sample type
NC
PC

**Figure S4.** Levels of IgM, IgA and IgG antibodies (median fluorescence intensity, MFI) to RBD in positive (TS) and negative samples (NC), and % seropositivity among TS, comparing 45 min versus 30 min incubation times of the secondary antibody conjugated to PE after 1 hour incubation of plasma samples with antigen beads.

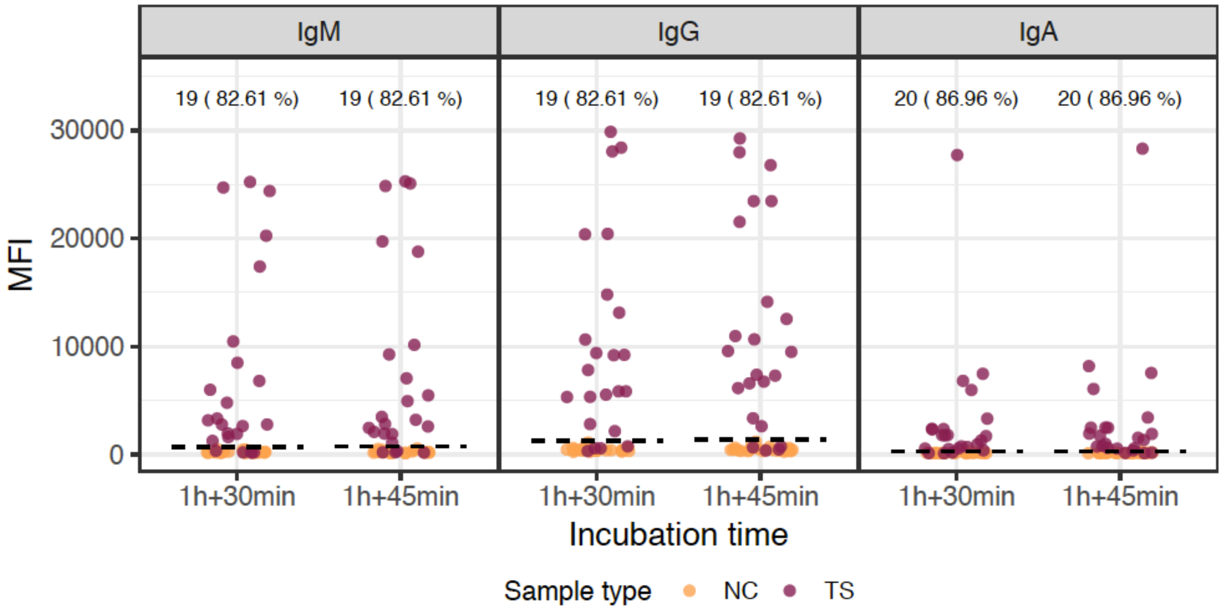

**Figure S5.** Ranking of isotype/antigen markers by Random Forest models for all negative controls plus either all positive samples or positive samples at different periods since onset of symptoms.

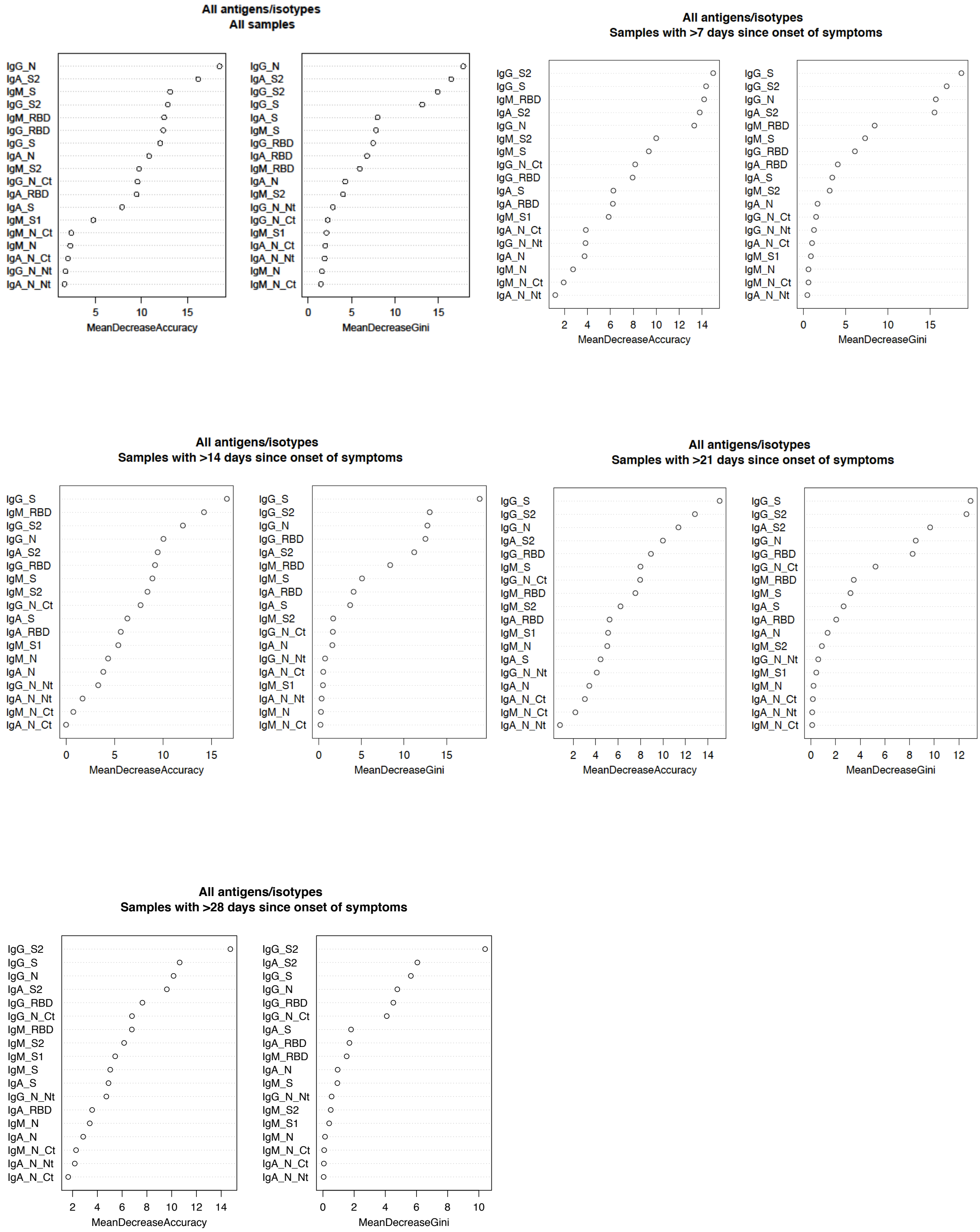

**SUPPLEMENTARY TABLES**

**Table S1.** Characteristics of individuals from whom positive samples were tested with regards to age, sex, symptoms, days since symptoms onset, and days since rRT-PCR diagnosis.

| **Continuous variable** | | **Median (IQR)** |
|---|---|---|
| **Age** | | 49.30 (26.24) |
| **Categorical variables** | **Category** | **N (%)** |
| **Sex** | Female | 71 (61.74) |
| | Male | 44 (38.26) |
| **Symptoms** | No | 8 (6.96) |
| | Yes | 107 (93.04) |
| **Hospitalized patients** | Yes | 49 (42.61) |
| **Days since onset of symptoms** | 0-6 | 13 (12.26) |
| | 7-13 | 14 (13.21) |
| | 14-20 | 28 (26.42) |
| | 21-27 | 24 (22.64) |
| | ≥28 | 27 (25.47) |
| **Days since first positive rRT-PCR** | 0-6 | 22 (19.13) |
| | 7-13 | 25 (21.74) |
| | 14-20 | 33 (28.70) |
| | ≥21 | 18 (15.64) |
| | After sample collection | 6 (5.22) |
| | Not available | 11 (9.56) |

**Table S2. Sensitivity and specificity of the assay** for negative controls plus either all positive samples or positive samples at different times since onset of symptoms and different thresholds targeting specificities of 100%, 99% and 98%.

| Antibody/antigen combinations | AUC | Specificity | Sensitivity |
|---|---|---|---|
| All samples | | | |
| IgA N + IgG N + IgG N Ct + IgM S2 | 0.918 | 100% | 82.61% |
| IgA N + IgA S + IgG N + IgM S2 | 0.917 | 100% | 82.61% |
| IgA S2 + IgG N + IgM S2 | 0.933 | 100% | 81.74% |
| IgA S2 + IgG N + IgM RBD + IgM S2 | 0.932 | 100% | 81.74% |
| IgA S2 + IgG N + IgG N Ct + IgM RBD + IgM S2 | 0.930 | 100% | 81.74% |
| IgA S2 + IgG N + IgG RBD + IgG S + IgG S2 + IgM RBD + IgM S | 0.930 | 99.22% | 83.48% |
| IgA S2 + IgG N + IgG S + IgG S2 + IgM RBD + IgM S | 0.930 | 99.22% | 83.48% |
| IgA S + IgA S2 + IgG N + IgG S + IgG S2 + IgM RBD | 0.923 | 99.22% | 83.48% |
| IgA S2 + IgG N + IgG S + IgG S2 | 0.920 | 99.22% | 83.48% |
| IgA S2 + IgG N + IgG RBD + IgG S + IgM RBD + IgM S | 0.935 | 99.22% | 82.61% |
| IgA S2 + IgG N Ct + IgM S2 | 0.934 | 98.44% | 83.48% |
| IgA S2 + IgG N + IgG RBD + IgG S + IgG S2 + IgM RBD + IgM S | 0.930 | 98.44% | 83.48% |
| IgA S2 + IgG N + IgG N Ct + IgG S + IgM RBD + IgM S2 | 0.930 | 98.44% | 83.48% |
| IgA S2 + IgG N + IgG S + IgG S2 + IgM RBD + IgM S | 0.930 | 98.44% | 83.48% |
| IgA S + IgA S2 + IgG N + IgG N Ct + IgG RBD + IgG S + IgM RBD + IgM S | 0.929 | 98.44% | 83.48% |
| ≥7 days since onset symptoms | | | |
| IgA S2 + IgG N + IgG S + IgM RBD + IgM S + IgM S2 | 0.975 | 100% | 92.47% |
| IgA S2 + IgG N Ct + IgM RBD | 0.973 | 100% | 92.47% |
| IgA RBD + IgA S2 + IgG N Ct + IgM RBD + IgM S2 | 0.972 | 100% | 92.47% |
| IgA RBD + IgA S2 + IgG N + IgG S2 + IgM RBD + IgM S2 | 0.971 | 100% | 92.47% |
| IgA S2 + IgM S | 0.971 | 100% | 92.47% |
| IgA S2 + IgG S2 + IgM RBD | 0.978 | 99.22% | 93.55% |
| IgA RBD + IgA S2 + IgG S2 | 0.976 | 99.22% | 93.55% |
| IgA RBD + IgA S + IgA S2 + IgG S2 + IgM RBD + IgM S | 0.974 | 99.22% | 93.55% |

| | | | |
|---|---|---|---|
| IgA S2 + IgG N Ct + IgG S2 + IgM RBD | 0.973 | 99.22% | 93.55% |
| IgA RBD + IgA S2 + IgG S2 + IgM RBD + IgM S2 | 0.973 | 99.22% | 93.55% |
| IgA S2 + IgG S + IgG S2 | 0.976 | 98.44% | 94.62% |
| IgA S2 + IgG S + IgM S | 0.986 | 98.44% | 93.55% |
| IgA RBD + IgA S2 + IgG S + IgM S | 0.984 | 98.44% | 93.55% |
| IgA RBD + IgA S2 + IgG S2 + IgM S | 0.983 | 98.44% | 93.55% |
| IgA S2 + IgG S2 + IgM S | 0.981 | 98.44% | 93.55% |
| ≥28 days since onset symptoms | | | |
| IgG N + IgG S + IgM S + IgM S2 | 0.998 | 100% | 96.30% |
| IgG N + IgG S + IgM RBD + IgM S2 | 0.998 | 100% | 96.30% |
| IgG N + IgG N Ct + IgM S2 | 0.978 | 100% | 96.30% |
| IgG N + IgG N Ct + IgM RBD + IgM S2 | 0.978 | 100% | 96.30% |
| IgA S2 + IgG N + IgM S2 | 0.976 | 100% | 96.30% |
| IgA S2 + IgG N + IgG S + IgM S2 | 0.999 | 99.22% | 96.30% |
| IgG N Ct + IgG S + IgM RBD + IgM S1 | 0.999 | 99.22% | 96.30% |
| IgA S2 + IgG RBD + IgG S + IgM S1 + IgM S2 | 0.999 | 99.22% | 96.30% |
| IgG N + IgG S + IgM RBD + IgM S1 + IgM S2 | 0.999 | 99.22% | 96.30% |
| IgG N Ct + IgG S + IgM RBD + IgM S1 + IgM S2 | 0.999 | 99.22% | 96.30% |
| IgA S2 + IgG RBD + IgG S + IgM S1 + IgM S2 | 0.999 | 98.44% | 100% |
| IgA S2 + IgG S + IgM S1 + IgM S2 | 0.998 | 98.44% | 100% |
| IgA S2 + IgG S + IgM S1 | 0.998 | 98.44% | 100% |
| IgG RBD + IgG S + IgM S1 + IgM S2 | 0.998 | 98.44% | 100% |
| IgA S + IgA S2 + IgG RBD + IgG S + IgM S | 0.998 | 98.44% | 100% |

### Table S3. Serological tests externally validated

| Test | Type | Antibodies | Manufacturer | Antigens | Positive controls | Negative controls | Specificity | Sensitivity | Reference |
|---|---|---|---|---|---|---|---|---|---|
| Biohit/Salocor RDT IgG/IgM | RDT | IgG/IgM | BIOHIT HealthCare Ltd | NP and RBD | 452 RT-PCR+ symptomatic | 1438 RT-PCR- | IgG 99.7-100%, IgM 97.6-100% | IgG/IgM 45.6-77.4% <14 days since onset symptoms; 94.1-100% ≥14days since onset symptoms | 1 |
| ABBOTT Architect SARS-CoV-2 IgG Assay | Chemiluminescent microparticle immunoassay | IgG | ABBOTT | NP | 125 RT-PCR+ symptomatic and hospitalized (689 samples) | 1020 pre-pandemia | 99.90% | 53.1% for 1-7 days since onset symptoms and 100% at day 17 since onset symptoms, and day 13 after PCR positivity | 2 |
| | | | | | 48 RT-PCR+ and symtomatic (103 samples) | 153: 80 patients RT-PCR-, 50 pre-pandemia, 5 with HKU1, NL63 or 229E, 4 with Influenza A or B, 14 with potentially interfering antibodies (CMV IgG, EBV VCA IgG and IgM, and rheumatoid factor) | 99.40% | 93.8% ≥14days since onset symptoms | 3 |
| | | | | | 71 RT-PCR+ symtomatic (113 samples) | 1182: 1063 pre-pandemic samples and 119 samples positive for antibodies against viruses and other pathogens to test cross-reactivity | 99.62% | 100% ≥ 14 days since onset symptoms | 4 |
| | | | | | 423 RT-PCR+ (no data on symptoms) | 1013 pre-pandemia | 99.80% | 29.3% at ≤7 days since onset of symptoms to 96.8% at ≥22 days since onset; overall sensitivity 80.4% | 5 |
| EUROIMMUN Anti-SARS-CoV-2 ELISA Assays for IgG | ELISA | IgG | EUROIMMUNE | S1 | 48 RT-PCR+ symtomatic (103 samples) | 153: 80 patients RT-PCR-, 50 pre-pandemia, 5 with HKU1, NL63 or 229E, 4 with Influenza A or B, 14 with potentially interfering antibodies (CMV IgG, EBV VCA IgG and IgM, and rheumatoid factor) | 94.80% | 94.8% ≥14days since onset symptoms | 3 |
| | | | | | 94 RT-PCR+ symptomatic (167 samples) | 103 pre-pandemia (49 respiratory infections, 14 cororavirus, 40 with antibodies to CMV, EBV, HIV) | 96.10% | 89.5% ≥14days since onset symptoms | 6 |
| | | | | | 181 RT-PCR+ (91 hospitalized patients and 90 from the outpatient clinic) | 176 pre-pandemia | 100% with optimized IgG ratio cut-off > 1.5 | 86% optimized IgG ratio cut-off > 1.5 | 7 |
| ROCHE Elecsys anti-SARS-CoV-2 | Electrochemiluminescent immunoassay (ECLIA) | Total antibodies (including IgG) | Roche Diagnostic | NP | 97 RT-PCR+ symtomatic (140 samples) | 79 pre-pandemia some with potential cross-reactive antibodies (antinuclear, antithyroglobulin, anti-Treponema pallidum, antistreptolysin O, anti-thyroid peroxidase, chikungunya, direct Coombs, hepatitis B, hepatitis C, hepatitis E, HIV, Chlamydia, Coxiella burneti, Borrelia, CMV, EBV, Mycoplasma pneumoniae, parvovirus B19, Toxoplasma gondii, influenza, RAI (search for irregular agglutinins), and rheumatoid factor) | 100% | 91.7% ≥14days since RT-PCR+ optimizing cut off with ROC 100%; 91.1% ≥14days since onset symptoms, optimizing cut off 95.1%; 96.7% ≥ 28 days since symptoms, optimizing cut off 100% | 8 |
| | | | | | 48 RT-PCR+ symtomatic (103 samples) | 153: 80 patients RT-PCR-, 50 pre-pandemia, 5 with HKU1, NL63 or 229E, 4 with Influenza A or B, 14 with potentially interfering antibodies (CMV IgG, EBV VCA IgG and IgM, and rheumatoid factor) | 98.69% | 89.36% ≥14 days since onset symptoms | 9 |
| Clungene COVID-19 IgG/IgM Rapid Test Cassette | RDT | IgG/IgM | Clongene Biotech, (Hangzhou,China) | no data | 94 RT-PCR+ symptomatic (167 samples) | 103 pre-pandemia (49 respiratory infections, 14 cororavirus, 40 with antibodies to CMV, EBV, HIV) | IgM 91.3%, IgG 98.1% | IgM 55.3%, IgG 97.4% ≥14days since onset symptoms | 6 |
| Orient Gene COVID-19 IgG/IgM Rapid Test cassette | RDT | IgG/IgM | Zheijang OrientGene Biotech, (Huzhou, China) | no data | 94 RT-PCR+ symptomatic (167 samples) | 103 pre-pandemia (49 respiratory infections, 14 cororavirus, 40 with antibodies to CMV, EBV, HIV) | IgM 95.1%, IgG 93.2% | IgM 97.4%, IgG 92.1% ≥14days since onset symptoms | 6 |
| VivaDiag™ COVID-19 IgM/IgG Rapid Test | RDT | IgG/IgM | VivaChek Biotech (Hangzhou, China) | no data | 94 RT-PCR+ symptomatic (167 samples) | 103 pre-pandemia (49 respiratory infections, 14 cororavirus, 40 with antibodies to CMV, EBV, HIV) | IgM 100%, IgG 99.0% | IgM 97.4%, IgG 94.7% ≥14days since onset symptoms | 6 |
| | | | | | 79 RT-PCR+ symptomatic (128 samples) | 108 pre-pandemia | IgM 94.9%, IgG 96% | IgM and IgG: 28% for 1–5 days since onset of symptoms, to 90% >20 days since onset of symptoms | 10 |
| StrongStep® COVID-19 IgG/IgM Combo Test | RDT | IgG/IgM | Liming Bio-Products (Jinangsu, China) | no data | 94 RT-PCR+ symptomatic (167 samples) | 103 pre-pandemia (49 respiratory infections, 14 cororavirus, 40 with antibodies to CMV, EBV, HIV) | IgM 99.0%, IgG 99.0% | IgM 50.0%, IgG 97.4% ≥14days since onset symptoms | 6 |
| Multi-G MGA 2019-nCoV IgG/IgM Rapid Test Cassette | RDT | IgG/IgM | Multi-G, (Antwerp, Belgium) made in china | NP | 94 RT-PCR+ symptomatic (167 samples) | 103 pre-pandemia (49 respiratory infections, 14 cororavirus, 40 with antibodies to CMV, EBV, HIV) | IgM 91.3%, IgG 97.1% | IgM 57.9%, IgG 97.4% ≥14days since onset symptoms | 6 |
| Prima COVID-19 IgG/IgM Rapid Test | RDT | IgG/IgM | Prima Lab SA (Balerna, Switzerland) | no data | 94 RT-PCR+ symptomatic (167 samples) | 103 pre-pandemia (49 respiratory infections, 14 cororavirus, 40 with antibodies to CMV, EBV, HIV) | IgM 93.2%, IgG 90.3% | IgM 68.4%, IgG 100% ≥14days since onset symptoms | 6 |
| Dynamiker 2019-nCOV IgG/IgM Rapid Test | RDT | IgG/IgM | Dynamiker Biotechnology (Tianjin, China) | NP | 94 RT-PCR+ symptomatic (167 samples) | 103 pre-pandemia (49 respiratory infections, 14 cororavirus, 40 with antibodies to CMV, EBV, HIV) | IgM 95.1%, IgG 99.0% | IgM 97.4%, IgG 94.7% ≥14days since onset symptoms | 6 |
| STANDARD Q IgM/IgG Duo | RDT | IgG/IgM | SD Biosensor, Gyeonggi-do, Korea | NP | 71 RT-PCR+ symtomatic (113 samples) | 1182: 1063 pre-pandemic samples and 119 samples positive for antibodies against viruses and other pathogens to test cross-reactivity | IgM 98.7%, IgG 100% | IgM 85.7%, IgG 100% | 4 |
| Wondfo Total Antibody Test | RDT | IgG+IgM | Wondfo, Guangzhou, China | no data | 71 RT-PCR+ symtomatic (113 samples) | 1182: 1063 pre-pandemic samples and 119 samples positive for antibodies against viruses and other pathogens to test cross-reactivity | 100% | 100% ≥ 14 days since onset symptoms | 4 |
| | | | | | 79 RT-PCR+ symptomatic (128 samples) | 108 pre-pandemia | 99.10% | 38.5% for 1–5 days since onset of symptoms, 89.5% for 16-20 days and 81.8% >20 days since onset of symptoms | 10 |

| | | | | | | | | | |
|---|---|---|---|---|---|---|---|---|---|
| COVID-PRESTO | RDT | IgG/IgM | AAZ-LMB (France) | no data | 133 RT-PCR+ symtomatic | 143 RT-PCR- symtomatic | 100% | 100% ≥ 15 days since onset symptoms | 11 |
| COVID-DUO | RDT | IgG/IgM | AAZ-LMB (France) | no data | 129 RT-PCR+ symtomatic | 143 RT-PCR- symtomatic | 100% | 100% ≥ 15 days since onset symptoms | 11 |
| LIAISON SARS-CoV-2 S1/S2 IgG test | Chemiluminescence immunoassay (CLIA) | IgG | DiaSorin | SP | 104 RT-PCR+ (210 samples): 31 patients hospitalized with moderate symptoms, 37 in the ICU with severe symptoms and 37 just RT-PCR+ | 1200: 1140 pre-pandemia, 60 RT-PCR- (10 of them infected by other coronaviruses (HCoV-229E, HCoV-HKU1, HCoV-OC43, or HCoV-untyped strain)) | 97-98.5% | 91.3% ≥ 5 days since diagnostic, 95.7% ≥ 15 days since diagnostic | 12 |
| EDI Novel Coronavirus COVID-19 IgM ELISA kit | ELISA | IgM | Epitope Diagnostics, United States | no data | 79 RT-PCR+ symptomatic (128 samples) | 108 pre-pandemia | 97.20% | 17.9 for 1–5 days since onset of symptoms, to 81.8 >20 days since onset of symptoms | 10 |
| EDI Novel Coronavirus COVID-19 IgG ELISA kit | ELISA | IgG | Epitope Diagnostics, United States | NP | 79 RT-PCR+ symptomatic (128 samples) | 108 pre-pandemia | 90.70% | 39.3 for 1–5 days since onset of symptoms, to 90.9 >20 days since onset of symptoms | 10 |
| COVID-19 IgM and IgG Rapid Test | RDT | IgG/IgM | BioMedomics Inc, Morrisville, NC, USA | RBD | 79 RT-PCR+ symptomatic (128 samples) | 108 pre-pandemia | IgM 87.9%, IgG 93.3% | IgM: 25.9% for 1–5 days since onset of symptoms, 84.2% for 16-20 days and 81.8% >20 days since onset . IgG: 22.2% for 1–5 days to 81.8% >20 days. | 10 |
| PerfectPOC Novel Corona Virus (SARSCoV-2) IgM/IgG Rapid Test Kit | RDT | IgG/IgM | Bioperfectus Technologies Co Ltd, Jiangsu, China | NP, SP | 79 RT-PCR+ symptomatic (128 samples) | 108 pre-pandemia | IgM 97.1%, IgG 98.1% | IgM: 39.9% for 1–5 days since onset of symptoms to 100% >20 days since onset . IgG: 25% for 1–5 days to 90% >20 days | 10 |
| Novel Coronavirus (SARS-CoV-2) IgM/IgG Combo Rapid Test-Cassette | RDT | IgG/IgM | Decombio Biotechnology Co Ltd, Beijing, China | no data | 79 RT-PCR+ symptomatic (128 samples) | 108 pre-pandemia | IgM 90.7%, igG 91.6% | IgM: 30.8% for 1–5 days since onset of symptoms to 90.9% >20 days since onset of symptoms. IgG: 26.9% for 1–5 days to 90.9% >20 days. | 10 |
| COVID-19 (SARSCoV-2) IgG/IgM Antibody Test Kit (Colloidal Gold) | RDT | IgG/IgM | DeepBlue Medical Technology Co Ltd, Anhui, China | no data | 79 RT-PCR+ symptomatic (128 samples) | 108 pre-pandemia | IgM 84.3%, IgG 99.1% | IgM: 42.9% for 1–5 days since onset of symptoms to 90.9% >20 days since onset of symptoms. IgG: 21.4% for 1–5 days to 81.8% >20 days. | 10 |
| Novel Coronavirus (2019-nCoV) Ab Test (Colloidal Gold) | RDT | IgG+IgM | Innovita Biological Technology Co Ltd, Qian'an, China | NP, SP | 79 RT-PCR+ symptomatic (128 samples) | 108 pre-pandemia | IgM 96.3%, IgG 100% | IgM: 14.8% for 1–5 days since onset of symptoms, 38.7% for 11-15 days and 16.7% >20 days since onset of symptoms. IgG: 25.9% for 1–5 days, 78.1% for 11-15 days and 66.7% >20 days. | 10 |
| COVID-19 IgG/IgM Rapid Test Cassette | RDT | IgG/IgM | Premier Biotech, Minneapolis, MN, USA | no data | 79 RT-PCR+ symptomatic (128 samples) | 108 pre-pandemia | IgM 98.1%, IgG 99.1% | IgM: 35.7% for 1–5 days since onset of symptoms to 90.9% >20 days since onset . IgG: 21.4% for 1–5 days to 81.8% >20 days. | 10 |
| SARS-CoV-2 IgM/IgG Antibody Rapid Test | RDT | IgG/IgM | Sure Biotech, New York, NY, USA; Wan Chai, Hong Kong | NP, SP | 79 RT-PCR+ symptomatic (128 samples) | 108 pre-pandemia | IgM 100%, IgG 100% | IgM: 10.7% for 1–5 days since onset of symptoms to 72.7% >20 days since onset . IgG: 17.9% for 1–5 days to 90.9% >20 days. | 10 |
| Coronavirus IgG/IgM Antibody (COVID-19) Test Cassette | RDT | IgG/IgM | UCP Biosciences, San Jose, CA, USA | no data | 79 RT-PCR+ symptomatic (128 samples) | 108 pre-pandemia | IgM 98.1%, IgG 98.1% | IgM: 25% for 1–5 days since onset of symptoms to 90.9% >20 days since onset . IgG: 25% for 1–5 days to 81.8% >20 days. | 10 |

CMV: cytomegalovirus; EBV: Epstein-Barr virus; HIV: human immunodeficiency virus; NP: Nucleocapsid protein; RDT: rapid diagnostic test, RT-PCR: Reverse transcription polymerase chain reaction; SP: Spike protein; VCA: virus capside antigen
